# Genetically engineering endothelial niche in human kidney organoids enables multilineage maturation, vascularization and de novo cell types

**DOI:** 10.1101/2023.05.30.542848

**Authors:** Joseph C. Maggiore, Ryan LeGraw, Aneta Przepiorski, Jeremy Velazquez, Christopher Chaney, Evan Streeter, Anne Silva-Barbosa, Jonathan Franks, Joshua Hislop, Alex Hill, Haojia Wu, Katherine Pfister, Sara E. Howden, Simon C. Watkins, Melissa Little, Benjamin D. Humphreys, Alan Watson, Donna B. Stolz, Samira Kiani, Alan J. Davidson, Thomas J. Carroll, Ondine Cleaver, Sunder Sims-Lucas, Mo R. Ebrahimkhani, Neil A. Hukriede

**Author notes:** Correspondence should be addressed to N.A.H. and M.R.E., Address for correspondence: 3501 5th Ave; 5061 BST3, Pittsburgh, PA 15213.

## Abstract

Vascularization plays a critical role in organ maturation and cell type development. Drug discovery, organ mimicry, and ultimately transplantation in a clinical setting thereby hinges on achieving robust vascularization of *in vitro* engineered organs. Here, focusing on human kidney organoids, we overcome this hurdle by combining an inducible *ETS translocation variant 2* (*ETV2*) human induced pluripotent stem cell (iPSC) line, which directs endothelial fate, with a non-transgenic iPSC line in suspension organoid culture. The resulting human kidney organoids show extensive vascularization by endothelial cells with an identity most closely related to endogenous kidney endothelia. Vascularized organoids also show increased maturation of nephron structures including more mature podocytes with improved marker expression, foot process interdigitation, an associated fenestrated endothelium, and the presence of renin^+^ cells. The creation of an engineered vascular niche capable of improving kidney organoid maturation and cell type complexity is a significant step forward in the path to clinical translation. Furthermore, this approach is orthogonal to native tissue differentiation paths, hence readily adaptable to other organoid systems and thus has the potential for a broad impact on basic and translational organoid studies.

**Translational Statement:** Developing therapies for patients with kidney diseases relies on a morphologically and physiologically representative *in vitro* model. Human kidney organoids are an attractive model to recapitulate kidney physiology, however, they are limited by the absence of a vascular network and mature cell populations. In this work, we have generated a genetically inducible endothelial niche that, when combined with an established kidney organoid protocol, induces the maturation of a robust endothelial cell network, induces a more mature podocyte population, and induces the emergence a functional renin population. This advance significantly increases the clinical relevance of human kidney organoids for etiological studies of kidney disease and future regenerative medicine strategies.

**Graphical Abstract:** Genetically engineered endothelial niche induces mature cell populations in human kidney organoids

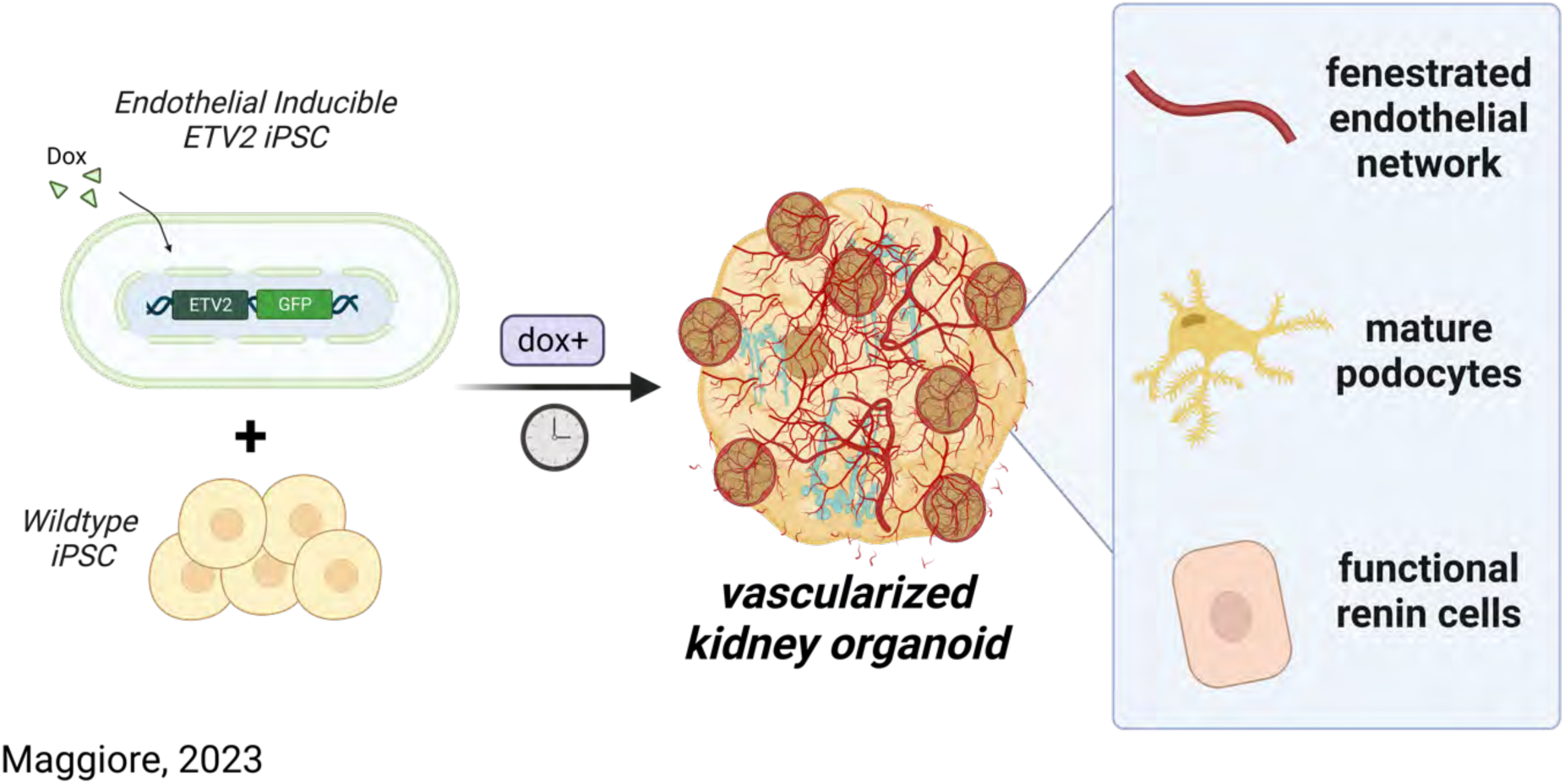

## Introduction

Engineering clinically relevant microphysiological tissues *in vitro* hinges on the co-development of a robust endothelial network. While endothelial cells play an important role in forming the vessels that transport blood and nutrients, they also interact with developing tissues and promote maturation^1–7^. Furthermore, across most organ systems, the vasculature plays a crucial role in disease processes. Therefore, *in vitro* organ models that lack the endothelial niche are limited in their capacity to recapitulate native physiology and model disease, and are also likely to be suboptimal in a transplant setting. Methods for vascularizing *in vitro* organ systems have been developed with varying levels of success, and have included bioprinting vasculature^8^, inducing vasculature using flow systems^9^, and supplementing cell culture systems with proteins or transcription factors^10^. Transgenic induction of an orthogonal hemato-endothelial population of cells^11–14^ is a particularly attractive approach given the hurdle it bypasses in developing a common universal culture media to coax the generation of heterogenous cell types. Overcoming this challenge, cells can be engineered with a transcriptionally guided endothelialization module that differentiates orthogonally to the parenchymal tissue of interest.

Recapitulating kidney vasculature in *in vitro* systems proves no less challenging. Kidney vasculature regulates blood pressure, controls filtration of serum ions and proteins, and regulates many important endocrine pathways^15^ and thus is critically important to model for studies of health and disease. Similarly, in disease settings, endothelial cells are known to fibrose and contribute to a patient’s disease burden in chronic kidney disease^16,17^. Current methods for vascularizing the *in vitro* human kidney^18^ have followed methods similar to other organ systems including flow enhancement with chips^19–21^, decellularizing/recellularizing ECM^22^, bioprinting^23^, renal subcapsular transplantation^24,25^, and VEGF supplementation^26^. However, these methods still leave multiple unmet needs^27^. Notably, there is a critical requirement for a method of vascularization that can: (1) reliably and simply generate a network that integrates with the kidney parenchyma, (2) allow for orthogonal differentiation and co-development of other non-endothelial cell types, (3) be broadly applied to multiple existing kidney tissue engineering protocols, (4) be highly reproducible, thereby minimizing batch to batch variation, and (5) be applicable in settings of high throughput kidney physiology and disease modelling.

For these reasons, we turned to *ETS translocation variant 2* (*ETV2*), previously shown to play a central role in directing endothelial cell differentiation^28,29^, to develop a genetically inducible hiPSC line (iETV2-hiPSC). We subsequently incorporated these cells into a previously established human kidney organoid bioreactor protocol^30,31^, to reconstitute natural endothelial niche in tandem with *in vitro* kidney organogenesis. The addition of the endothelial niche improves the maturation of podocytes with signs of vascular invasion and fenestration, induces the formation of a population of renin-positive cells, and results in significant cell-cell interactions between endothelial and parenchymal cells.

## Results

### ETV2 inducible hiPSCs generate endothelial cells following doxycycline exposure

We developed a doxycycline inducible ETV2-EGFP hiPSC line (Supplementary Fig. 1) that, when stimulated, generates a population of endothelial progenitor-like cells (Fig. 1a). In monolayer iPSC culture, following 24-hour doxycycline exposure there was a marked induction in immunofluorescence of ETV2-EGFP that persisted until day 2. Beginning at day 3, with subsequent continued doxycycline exposure, there was an increase in immunofluorescence intensity until day 6 of MCAM and PECAM1 expression (Fig. 1b-c, Supplementary Fig. 2-3). ERG, a transcription factor critical to endothelial cell differentiation and function^32^, was expressed on immunofluorescence at day 4 (Supplementary Fig. 4). Subsequently, MCAM and PECAM1 reach peak protein expression levels by day 7, which was confirmed by per cell fluorescence intensity quantification (Fig. 1b-c).

**Fig. 1:**
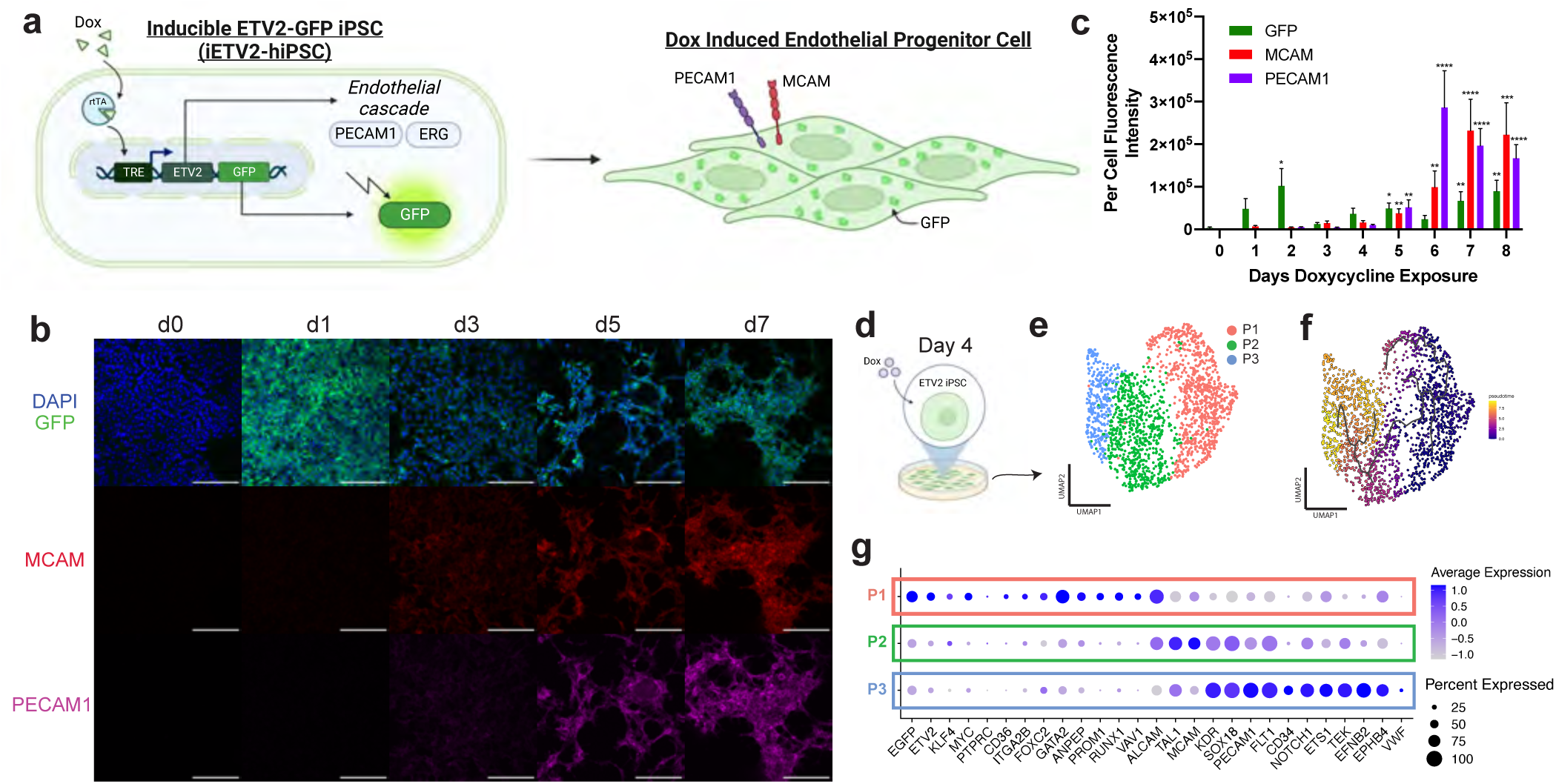
Genetically engineered iETV2-hiPSCs undergo synthetic endothelial differentiation. **(a)** Schematic of iETV2-hiPSC genetic circuit. **(b)** iETV2-hiPSCs differentiated with doxycycline induction and immunofluorescence from day 0 to 9; scale bar = 200μm. **(c)** Per cell immunofluorescent intensity of GFP, MCAM and PECAM1. N = 3 biological replicates; p-value **** : < 0.0001, *** : < 0.001, ** : < 0.01, * : < 0.05. **(d)** scRNAseq was carried out on iETV2-hiPSCs exposed to doxycycline for 4 days. **(e)** UMAP generated three distinct cellular populations. **(f)** Pseudotime trajectory analysis with Monocle3 demonstrated developmental lineage from P1 -> P2 -> P3. **(g)** Differentially expressed genes for P1, P2, P3.

**Fig. 2:**
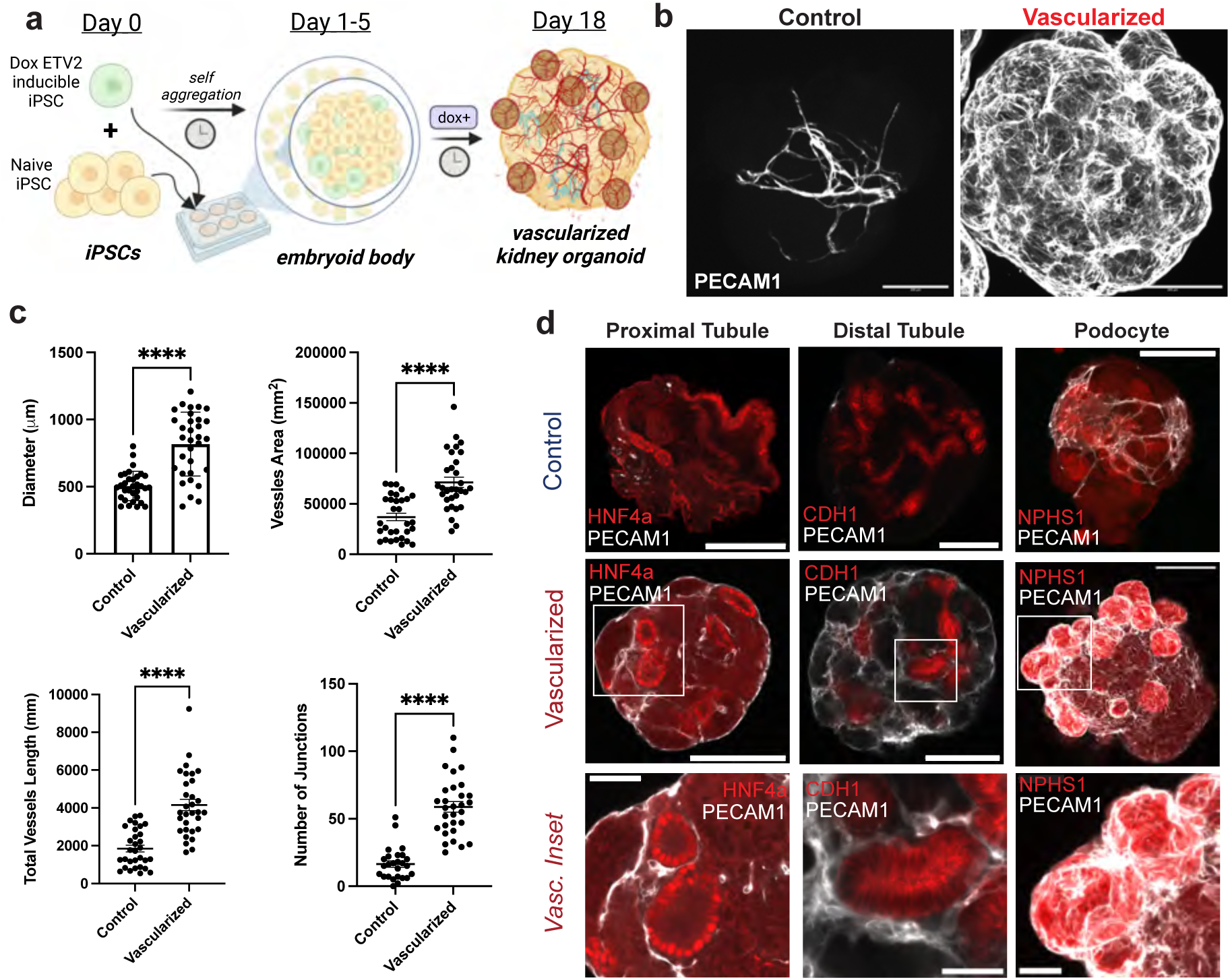
Vascularization of human kidney organoid. **(a)** Method for vascularizing human kidney organoids by combining iETV2-hiPSCs with wildtype iPSCs. **(b)** Immunofluorescence of endothelial cell network between MANZ2-2 control and vascularized kidney organoids from 3 independent biological replicates, showing representative images. **(c)** Angiotools quantification and diameter measurement between control and vascularized kidney organoids; organoids originate from 3 biological replicates. p-value **** : < 0.0001. **(d)** Representative immunofluorescence images of endothelial interaction with podocytes, proximal tubule, distal tubule, and stroma between control and vascularized kidney organoid. Organoid images scale bar = 200μm, inset scale bar = 50μm.

**Fig. 3:**
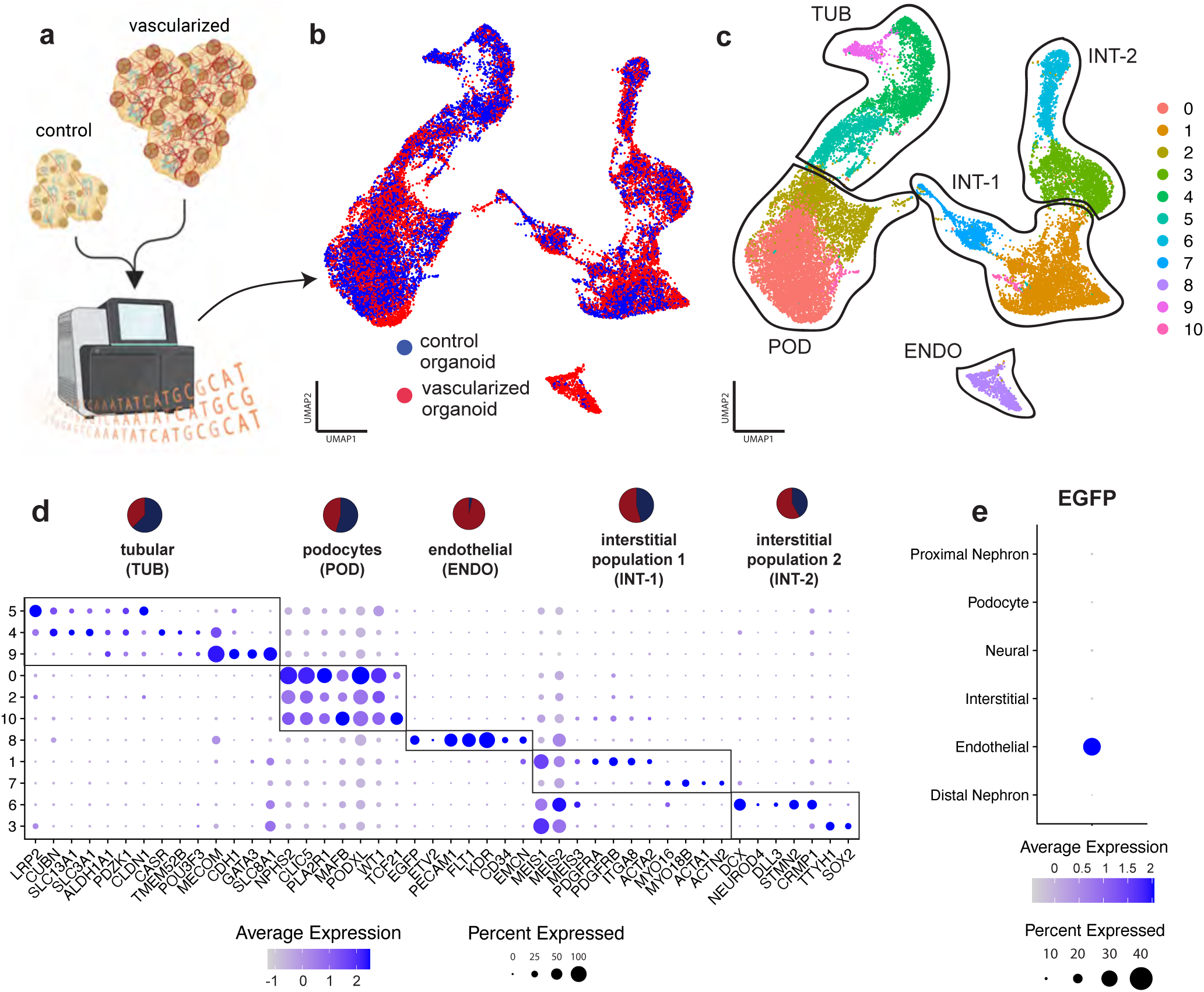
snRNAseq of control and vascularized human kidney organoids. **(a)** MANZ2-2 control and vascularized human kidney organoids were analyzed via snRNAseq. **(b)** Cells aggregated and well overlapped with Harmony, and did not segregate by batch. **(c)** Cells clustered by cell type as podocyte (POD), endothelial (ENDO), tubular (TUB), and interstitial (INT-1/2). **(d)** Differentially expressed genes per cluster were identified. Cellular populations were quantified to analyze composition by control or vascularization origin and graphed as pie chart per cell type (red = vascularized, blue = control). **(e)** EGFP expression was localized to endothelial population.

To determine the transcriptomic identify of the induced endothelial progenitor-like population, scRNAseq was performed on iETV2-hiPSCs exposed to doxycycline for 4 days (Fig. 1d). 1,839 cells were used in downstream scRNAseq analysis and UMAP plots were generated wherein three distinct cell clusters were identified (P1, P2, P3) (Fig. 1e). As identified using Monocle3^33,34^, a developmental trajectory was identified progressing from P1 -> P2 -> P3 (Fig. 1f). These clusters tended to contain transcriptional profiles similar to early, mid, and late endothelial progenitor populations (Fig. 1g). The early endothelial-like progenitor population (P1) was characterized by high levels of *ETV2*, *KLF4*, *GATA2*, and *RUNX1*; mid endothelial-like progenitor (P2) cells were characterized by *ALCAM*, *TAL1*, *MCAM*; and late endothelial-like progenitor (P3) were characterized by *PECAM1*, *FLT1*, and *CD34* (Fig. 1g, Supplementary Fig. 5).

To understand whether the induced iETV2-hiPSC endothelial cells differentiate into an organ-specific or endothelial sublineage, the single cell transcriptomes of P1, P2 and P3 populations were compared with published human endothelial scRNAseq datasets from Tabula Sapiens^35^ using SingleCellNet^36^. We first determined that the iETV2-hiPSC endothelial cells did not specifically align or express markers of organ-specific endothelial cell types (Supplementary Fig. S6-8). Furthermore, these cells do not express high levels of *EFNB2* or *EPHB4*, which mark arteries and veins, respectively (Supplementary Fig. 9). Interestingly, when the iETV2-hiPSCs were aligned against Tabula Sapiens endothelial cells gene ontology classifiers, there was increased alignment with terms indicative of endothelial cell vascular tree morphology (Supplementary Fig. S8). Taken together, this data demonstrated that when cultured in monolayer, iETV2-hiPSCs generate a generic, organ-agnostic population of endothelial-like progenitor cells.

### Kidney organoid vascularization method generates integrated endothelial network with maintained presence of critical kidney cell types

We modified an established kidney organoid protocol^31,37,38^ to include the integration of iETV2-hiPSCs (Fig. 2a). Vascularized human kidney organoids were generated by combining a wildtype hiPSC (MANZ2-2 or Triple) line with the iETV2-hiPSC transgenic line. Both hiPSCs were scraped from monolayer culture, dissociated as clusters in suspension, and when combined, self-aggregated into spheroids, forming embryoid bodies by day 3. From culture day 5 to 18, the organoids were exposed to doxycycline, thereby inducing the transgenic iETV2-hiPSC line to differentiate into endothelial cells. The protocol was optimized for minimum effective doxycycline concentration, doxycycline exposure window, doxycycline length and transgenic hiPSC to wildtype ratio to yield an extensive vascular network without compromising kidney organoid parenchymal cell types (Supplementary Fig. 10-11). Independent of changing the ratio of cell types, there was a consistent number of EGFP+PECAM1+ cells that form, indicating a potential self-regulation of endothelial cell number (Supplementary Fig. 12). We found that the ideal protocol to yield vascularized kidney organoids was a combination of 1:5 ETV2-EGFP hiPSCs : wildtype hiPSCs, induced with 0.5μg/mL doxycycline from day 5-18 (Supplementary Fig. 10-12).

By day 18, a significant endothelial network formed throughout the entire kidney organoid as demonstrated by PECAM1 (Fig. 2b), endomucin and NRP1 immunofluorescence (Supplementary Fig. 13) and by RT-qPCR (Supplementary Fig. 14). The overall increase in endothelial network with this vascularization method was analyzed with Angiotool^39^ on immunofluorescent PECAM1 images showing ∼2-fold increase in vessel area (p-value < 0.0001), ∼2-fold increase in total vessel length (p-value < 0.0001), and ∼3.60-fold increase in the number of junctions (p-value < 0.0001) (Fig. 2c). Although the total number of starting cells was the same between the MANZ2-2 control and vascularized organoids, the diameter of vascularized organoids was found to be ∼60% larger than the control kidney organoid (p-value < 0.0001) (Fig. 2c). To determine the functionality and stability of vessels generated within the organoids, we assessed if they could connect with an *in vivo* blood supply and become patent. To achieve this, we inserted human kidney organoids with or without iETV2-hiPSCs under the kidney capsule of nod-SCID gamma (NSG) mice as previously described^24^. These organoids were allowed to mature *in vivo* for 28 days. Vessels in organoids that did not contain the iETV2-hiPSCs contained vessels that were entirely derived from the mouse as previously shown^24^. Conversely, the iETV2-hiPSCs vascularized kidney organoids contained vessels that stained positive for human GAPDH and connected with the rodent vessels at the periphery of the organoid (Supplemental Fig. 15). Overall, this data demonstrated that the addition of iETV2-hiPSCs generates a robust endothelial network within the kidney organoid that can be sustained with *in vivo* connection with mouse vasculature.

Furthermore, with the vascularized kidney organoid protocol, there appeared to be interactions between other key epithelial cell types and the endothelial cells. Large nodules of cells were evident at the cortex of the vascularized kidney organoids, which appeared to be largely composed of NPHS1+ cells, a podocyte slit diaphragm specific marker^40^, enveloped by endothelial cells (Fig. 2c). Additionally, proximal, and distal tubules, imaged in cross-section, appear to be encased in endothelial cells, which was not observed in the MANZ2-2 control kidney organoid (Fig. 2c). However, other than the gross changes in organoid morphology, and distinct cell-cell interactions between epithelial-endothelial cells, we did not notice differences in the cell type morphology of epithelial cells, indicating preservation of critical cell type morphology (Fig. 2c).

To further understand the organoid cellular composition changes upon vascularization, snRNAseq was carried out on day 18 MANZ2-2 control and vascularized kidney organoids (Fig. 3a). On UMAP, cells from control and vascularized kidney organoid (Fig. 3b) clustered by distinct cell types – podocytes, proximal nephron, distal nephron, endothelial cells, and interstitial cells (Fig. 3c), and express characteristic cell-type specific markers (Fig. 3d). These cellular population annotations were confirmed using DevKidCC^41^ (Supplementary Fig. 16). The endothelial population (966 cells) was comprised almost entirely from endothelial cells of the vascularized kidney organoid (888 cells, 91.9%) and nearly all *EGFP* expression was localized to the endothelial population (Fig. 3e). The total organoid endothelial cell percentage for the control kidney organoid was 0.61% compared to 6.9% under vascularized conditions. Podocyte (POD) and interstitial cluster 1 (INT-1) populations contained evenly distributed populations between control and vascularized kidney organoid cells (54.4% and 45.6%, respectively) (Fig. 3d). Tubular segments contained a slightly higher percentage of cells from the control kidney organoid (62.3%). Interstitial Cluster 2 was comprised of cells in slightly greater proporitions from the vascularized kidney organoid (58.0%) (Fig. 3d). Tubular segments subclustered into proximal and distal tubule did not show significant expression differences between the control and vascularized conditions (Supplementary Fig. S17). Based on the initial cell type clustering and immunofluorescence analysis, we determined the vascularized kidney organoids contain a significant endothelial population while preserving critical epithelial cell types. Next, we investigated the distinct cellular changes due to the addition of the iETV2-hiPSCs that occurred in the podocyte population.

### Vascularized kidney organoid enables greater differentiation of podocyte populations and improved glomerular vascular interaction

Podocytes are a critical cell type in kidney physiology, acting as one of the early regulators of blood filtration. They have a highly specialized morphology consisting of interdigitating foot processes that encase endothelial cells with glomerular basement membrane (GBM) in between^42^. Modelling of these foot processes with GBM is necessary to adequately understand mechanisms of glomerular filtration or identify therapeutics for diseases such as diabetic nephropathy, focal segmental glomerulosclerosis, and lupus nephritis^43,44^. As previously reported, podocyte-like cells are present in MANZ2-2 control organoids, however, they generally have little to no endothelial cell integration, foot process development, or glomerular basement membrane^27,45^ (Fig. 4a-c). Other studies have examined podocyte-endothelial interaction and have found present but limited vascular involvement^46^. Podocytes in MANZ2-2 control kidney organoids exist as a defined cluster of cells on the exterior surface of the kidney organoid as NPHS1+ by immunofluorescence, with substantiative microvilli projections on scanning electron microscopy (Fig. 4a, c). The podocytes contain numerous protrusions, possibly representing primitive foot processes, which are disconnected from endothelia, and on transmission electron microscopy are seen to extend out from the exterior surface randomly or contact similar protrusions from other podocytes (Fig. 4c). By contrast, vascularized human kidney organoids contain an extensive EGFP+ PECAM1+ endothelial cell network that both encapsulates and invaginates into the podocyte cluster (Fig. 4b). Furthermore, on transmission electron microscopy, we detect identifiable interactions of interdigitating foot processes. (Fig. 4c). Our data demonstrates that the presence of iETV2-hiPSCs endothelial cells enables direct cell-cell interactions with the podocytes, thereby enabling greater physiologically relevant morphological features.

**Fig. 4:**
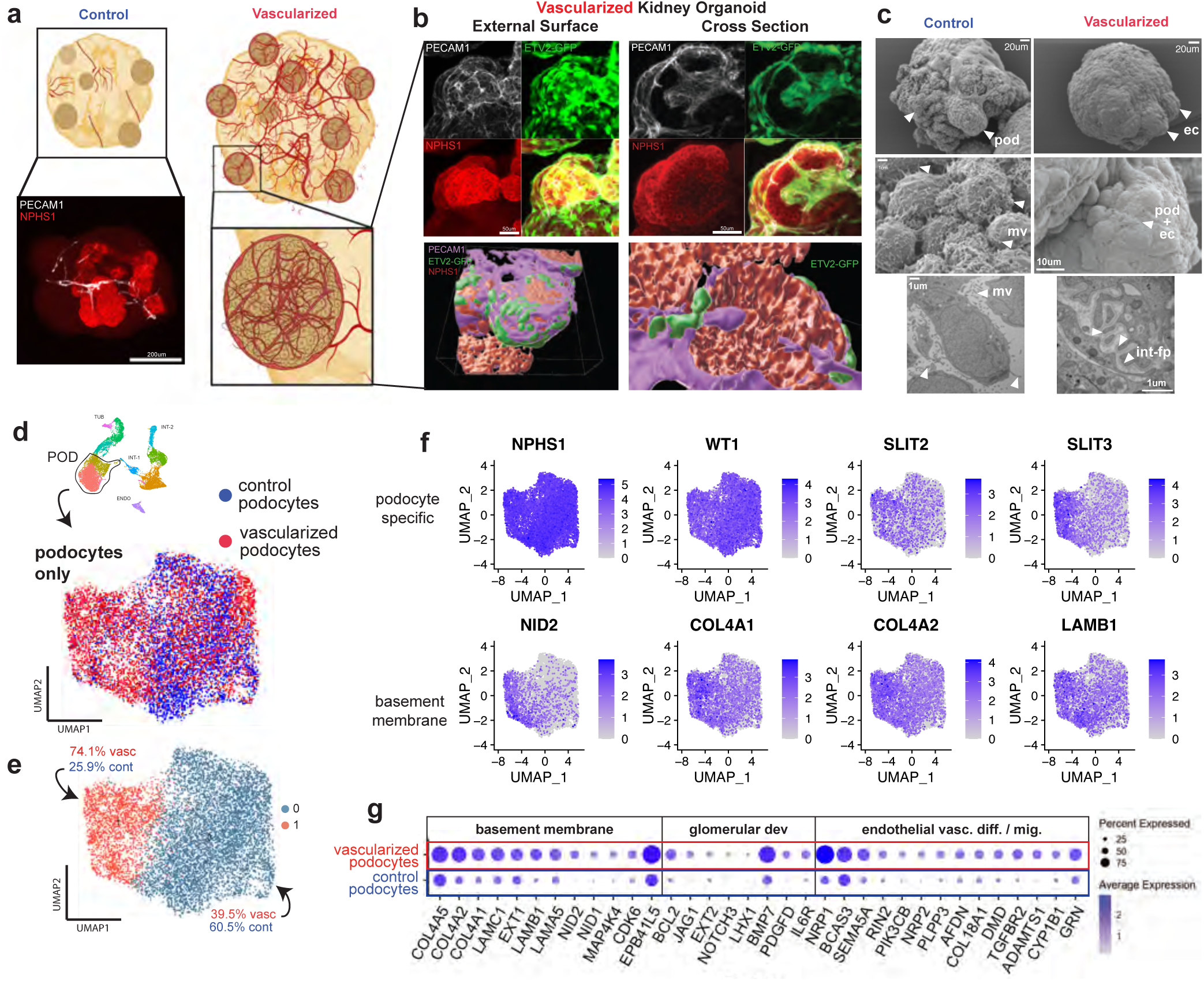
Increased maturation of podocytes with vascularization. **(a)** Podocytes exist in clusters in kidney organoids on the exterior surface; they become highly vascularized with the vascularization protocol and lack vascular integration with the control kidney organoid. **(b)** Vascularized MANZ2-2 kidney organoids contain GFP+ endothelial cells encasing the podocyte clusters from the external surface, and invaginating networks through the middle of the cluster. **(c)** SEM and TEM of control and vascularized kidney organoid. pod: podocyte; ec: endothelial cell; fp: foot process; fen: fenestration; gbm: glomerular basement membrane; int-fp: interdigitating foot processes. **(d)** UMAP of podocytes from both control and vascularized kidney organoid **(e)** Podocytes distinctly cluster into two populations largely predominated by control or vascularized podocytes. **(f)** Podocyte specific markers are present in both clusters, however slit diaphragm and basement membrane markers are upregulated in Cluster 0, predominated by vascularized podocytes. **(g)** Gene set enrichment analysis identifies upregulated pathways of basement membrane, glomerular development and endothelial vasculature differentiation and migration in the vascularized predominating podocyte cluster.

To better interrogate podocyte maturation, snRNAseq podocytes from control and vascularized kidney organoids were further separated into two distinct clusters on UMAP (Fig. 4d). Cluster 1 was largely comprised of podocytes from the vascularized kidney organoid (74.1%), while Cluster 0 was largely comprised of podocytes from the control kidney organoid (60.5%) (Fig. 4e). Prior studies have investigated the development of podocytes through human fetal kidney transcriptomics and found two populations of podocytes with distinct gene expression patterns – early podocytes (*OLFM3, GFRA3, PAX8, LHX1, PCDH9*) and late podocytes (*CLIC5, PLCE1, PTPRO, NPHS1, NPHS2*)^47,48^. We find that podocytes in MANZ2-2 control, and vascularized kidney organoids contain higher levels of late markers than early markers (Supplementary Fig 18a).

Interestingly, podocytes of the vascularized kidney organoid contain higher expression levels for markers of slit diaphragm and basement membrane formation, such as *SLIT2, SLIT3, NID2, COL4A1, COL4A2, and LAMB1* (Fig. 4f). Furthermore, a gene ontology search was conducted for the top 100 differential expressed genes from the vascularized podocyte predominant cluster, which resulted in marked increase in gene ontology matches for basement membrane formation, glomerular development, endothelial vascular differentiation and migration, and tight junction regulation (Fig. 4g, Supplementary Fig. 19). Previous studies have described the maturation of podocyte basement membrane composition by a transition of COL4A1/2 to COL4A3/4/5 and a transition in laminins from LAMA1 to LAMC1 to LAMA5^44,49^. We found that podocytes of the vascularized kidney organoid contained higher proportions of mature basement markers than the control podocytes, such as *COL4A5*, *LAMC1* and *LAMA5* (Supplementary Fig. 18b). Finally, using CellChat, it was determined that podocytes from the vascularized predominant cluster had significantly higher *VEGF* signaling as compared with the control podocytes (Supplementary Fig. 18c). Taken together, this data demonstrates that the iETV2-hiPSCs endothelial niche within the vascularized kidney organoid promotes the emergence of a more mature podocyte population. We next investigated the effect of added iETV2-hiPSCs on the interstitium of the vascularized kidney organoid.

### iETV2-hiPSCs niche in vascularized organoid enables generation of a renin positive cellular population in interstitium

The interstitial population of the kidney is a heterogenous population^50–52^ that importantly contributes to the development, maturation, and structural support of other kidney parenchymal cell types^53–55^. To further investigate the interstitial cell populations identified from the snRNAseq analysis, INT-1 cell population was re-clustered on UMAP (Fig. 5a). INT-1 was found to contain 7 subclusters, comprised of vascular smooth muscle renin+ cells (VSM-REN), two fibroblast populations (FIB-1 & FIB-2), proliferating fibroblast cells (PROL-FIB), mural cells (MUR), myofibroblasts (MYO), and proliferating myofibroblasts (PROL-MYO) (Fig. 5b). Cell type specific markers aligned with previously published work identifying interstitial heterogeneity^56,57^ (Fig. 5c, Supplementary Fig. 20-21).

**Fig. 5:**
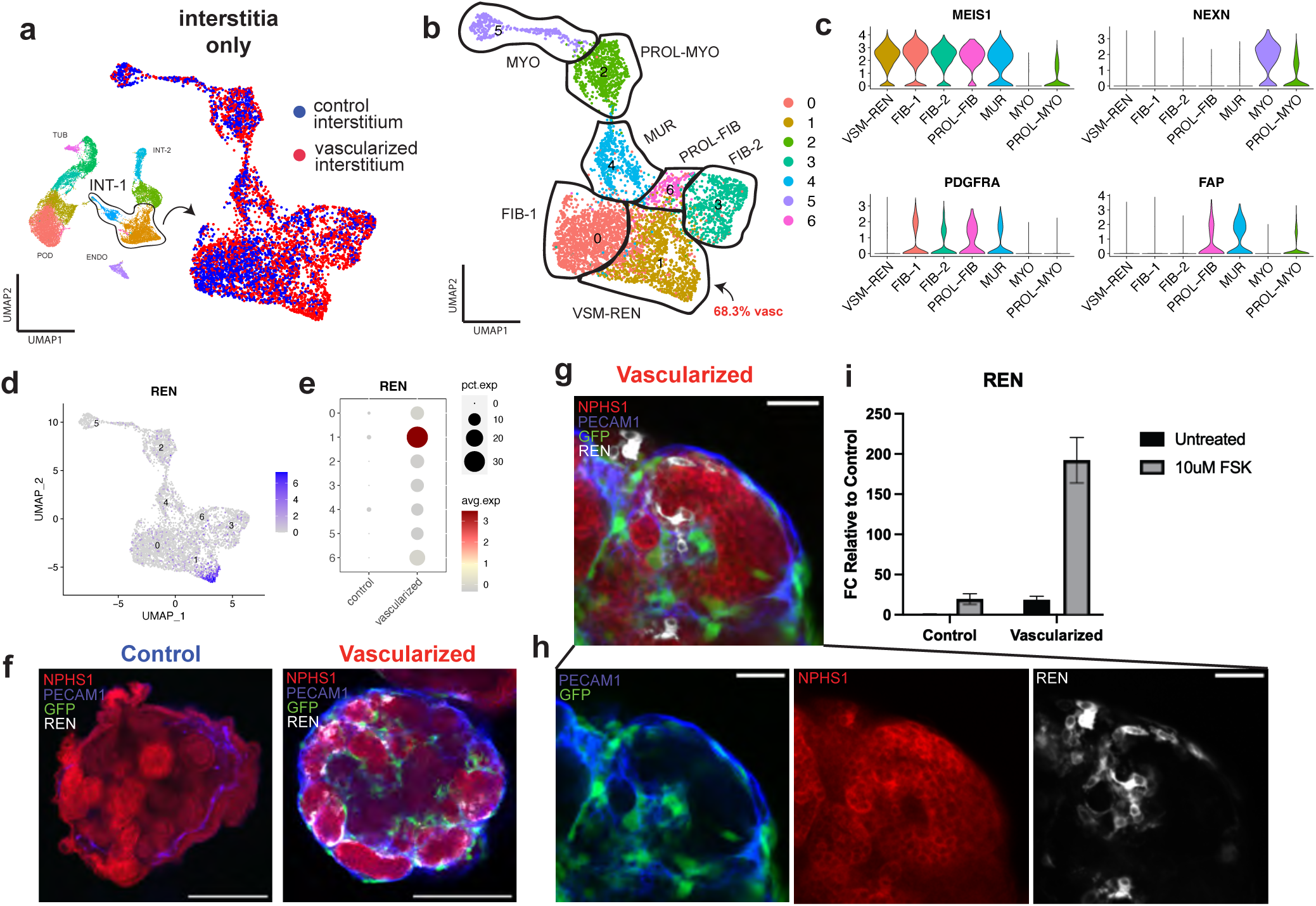
Vascularization of kidney organoid enables emergence of a renin cell population. **(a)** UMAP of interstitial cells from MANZ2-2 control and vascularized kidney organoid. **(b)** Interstitial cells cluster into 8 distinct populations. MUS: muscle; MUR: mural; INT-X: interstitial population-X; REN: renin cell; ABB-POD-FIB/ABB: abberant podocyte-fibroblast population. **(c)** Violin plots of key genes per cluster. **(d)** REN specifically localized to a population of cells that **(e)** largely originates from the vascularized kidney organoid. **(f)** Control kidney organoid contains no REN+ cells on immunofluorescence while vascularized kidney organoid contains many spread across podocyte (NPHS1+) clusters; scale bar = 200μm. **(g)** Vascularized kidney organoid podocyte clusters contain REN+ cells within the cluster, **(h)** juxtaposing but not colabelling with NPHS1+ podocytes or GFP+PECAM1+ endothelial cells; scale bar = 50μm. **(i)** 10uM forskolin (FSK) – a pro renin stimulatory drug – on organoids enables 200-fold increase in renin expression in the vascularized kidney organoid.

Contained within the *in vivo* kidney’s interstitial population is a group of cells known as the juxtaglomerular apparatus, which contain renin positive cells that regulate blood pressure through the renin-angiotensin system (RAS)^58^. Renin cells, when diseased, contribute to hypertension, chronic kidney disease, and diabetic nephropathy^59,60^. Therefore, a representative *in vitro* recapitulation of this cellular population is highly needed and renin production remains an important physiological assay^61^. Wildtype organoids have not been found to contain a resident renin expressing population. Only when stimulated with forskolin do kidney organoids produce renin^62,63^. We identified for the first time, the emergence of renin cells in the vascularized kidney organoid without using exogenous stimuli such as forskolin (Fig. 5d-e). REN+ cells were confirmed by immunofluorescence in the vascularized kidney organoids (Fig. 5f-h). These REN+ cells were localized to the podocyte clusters, whereby they do not co-label as PECAM1+ or NPHS1+ indicating that they may be forming a portion of a rudimentary juxtaglomerular apparatus^64,65^. Furthermore, REN+ cells in the vascularized kidney organoid are EGFP- (Supplementary Fig. 2) and REN+ cells do not exist in the MANZ2-2 control kidney organoid. We therefore demonstrate that the existence of the iETV2-hiPSC endothelial niche enables the formation of a population of renin cells without the usage of exogenous stimulation.

Drugs such as forskolin, can be used as a functional assay to trigger the release of renin in kidney organoid stromal cells^62,63^. We demonstrated that our vascularized kidney organoids at a basal level express a similar level of renin as the control MANZ2-2 kidney organoids that are stimulated with 10μM forskolin. When the vascularized kidney organoids are stimulated with 10μM forskolin, the organoids express 192.4-fold increase in renin expression relative to MANZ2-2 control unstimulated kidney organoids and 9.8-fold increase in renin expression relative to unstimulated vascularized kidney organoids (Fig. 5i). Other studies have shown that forskolin also triggers tubular swelling in control organoids through cAMP activation^40^. We observed the cyst formation with forskolin stimulation to be larger in the vascularized kidney organoid than the control kidney organoid (Supplementary 27) indicating that the iETV2-hiPSCs endothelial niche may enable greater electrophysiological ion transport than the control organoid^61^. This data taken together, demonstrates for the first time that the iETV2-hiPSCs endothelial niche induces, in the vascularized kidney organoid, a functional renin population without the usage of exogenous stimulation. This serves as critical step forward in microphysiological modelling of systems for blood pressure mechanistic understanding and drug discovery. Finally, we aimed to investigate whether the iETV2-hiPSCs added to the kidney organoid co-develop with the maturing epithelial cells of the vascularized kidney organoid.

### Co-developing iETV2-hiPSCs produce kidney specific endothelial network within kidney organoids

Previous studies across many organ systems have found that there is a significant crosstalk between endothelial and epithelial cell types – positing that the crosstalk is necessary to inform proper co-development and maturation between one another^1,7,45,66^. To understand whether iETV2-hiPSCs mature and adopt an organ-specific fate, we reclustered the snRNAseq endothelial cell populations from control and vascularized kidney organoid for further analysis (Fig. 6a). The endothelial population was comprised of two subclusters, despite consistent canonical endothelial marker expression across all cells (PECAM1+ CDH5+) (Fig. 6b-c). Cluster 1 was found to be largely comprised of less mature endothelial cells indicated by expression of *ETV2* and *TAL1* while Cluster 2 was comprised of more differentiated endothelial cells indicated by expression of *PECAM1*, *EMCN*, *CLDN5*, and *TIE1* (Fig. 6d-f). The vascularized kidney organoid cells were then compared to iETV2-hiPSC induced endothelial cells grown in monolayer cultures (Fig. 1e). This analysis revealed that Cluster 1 from the organoids (EC1) aligned closest with the P2 cluster (53.0%), followed by the P3 cluster (39.2%), while Cluster 2 from the organoids endothelia aligned closest with the P3 cluster (47.3%), followed by the P2 cluster (35.1%) (Supplementary Fig. 24). Despite progression in maturity, the organoid endothelial cells do not show high levels of either *EFNB2* (artery) or *EPHB4* (vein) expression (Supplementary Fig. 25). Based on these data, we conclude that the iETV2-hiPSCs mature within the kidney organoid, progressing to a more differentiated endothelial state, but may retain adaptability and plasticity given their limited progression to either an artery or vein identity.

**Fig. 6:**
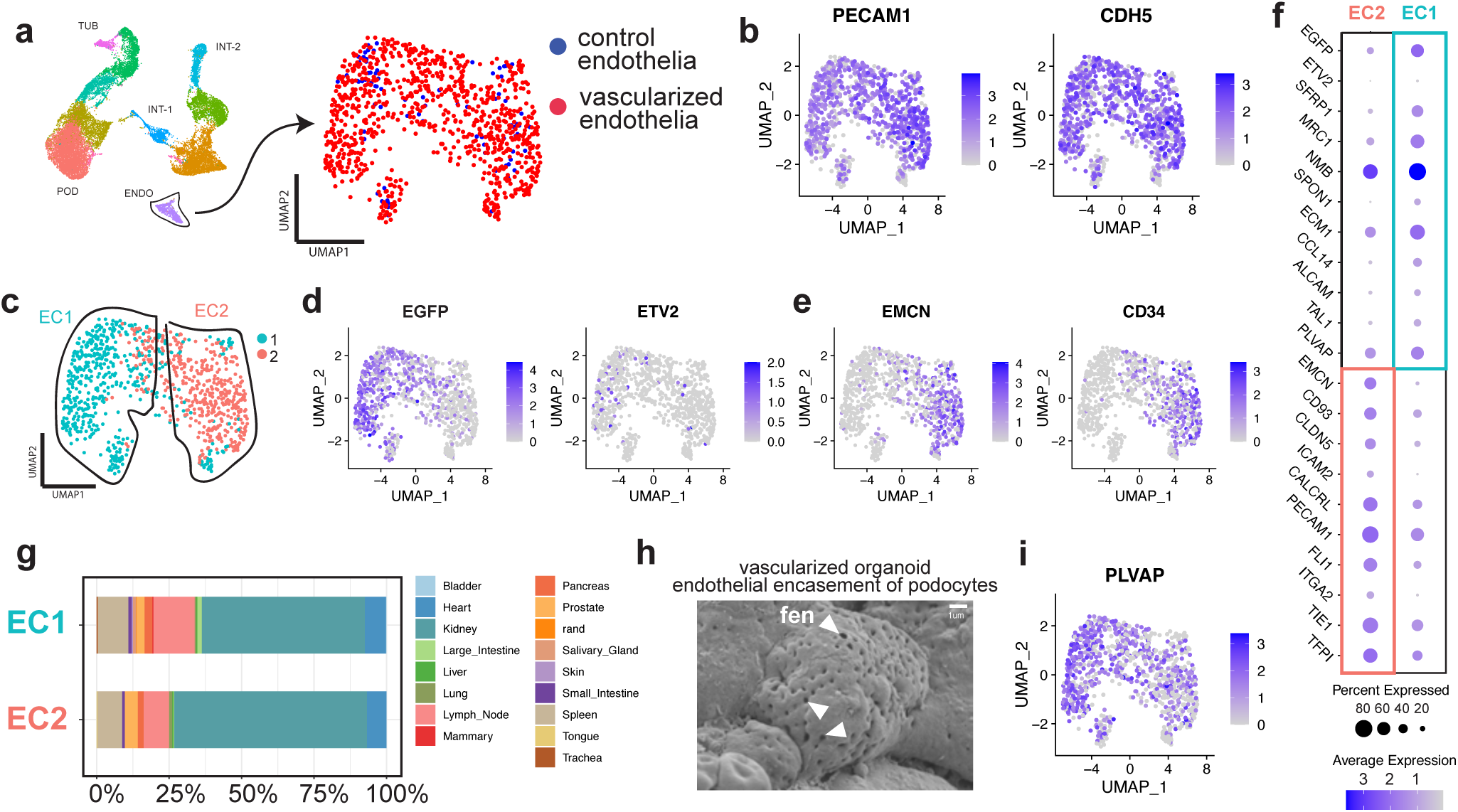
iETV2-hiPSCs undergo maturation and organ specification. **(a)** Endothelial population reclustered on UMAP. **(b)** Endothelial cells are PECAM1+ CDH5+. **(c)** Endothelial population cluster into two distinct populations consisting of **(d)** EC1 predominated by EGFP and **(e)** EC2 predominated by greater endothelial maturation markers EMCN and CD34. **(f)** Differentially expressed genes between EC1 and EC2. **(g)** SingleCellNet classification of EC1 and EC2 using Tabula Sapiens organ specific endothelial dataset demonstrates kidney specification of endothelial cells. **(h)** Vascularized human kidney organoids contain podocyte clusters encased by fenestrated endothelia. **(i)** iETV2-hiPSC derived endothelia express fenestration marker *PLVAP*.

To determine if the observed increase in maturity is accompanied by kidney-specific endothelial maturation, the endothelial cells of the kidney organoid were compared to organ-specific endothelia of Tabula Sapiens using SingleCellNet. We found that the iETV2-hiPSC population of the vascularized kidney organoid gained a kidney specific endothelial profile when incorporated into the kidney organoid protocol in contrast to the organ-non-specificity of the original iETV2-hiPSC monolayer. Cluster 1 and 2 endothelial cells from the organoid align as 56.2% and 65.8% kidney specificity, respectively, and 13.7% and 9.6% lymph node specificity, respectively (Fig. 6g). This data demonstrates that the *ETV2*-induced organoid endothelial cells adopt an organ-specific maturity in co-development with the kidney organoid. We also found that the maturation of iETV2-hiPSCs in vascularized kidney organoids also was accompanied by changes in transcriptional markers of vascular morphology, as was shown in the iETV2-hiPSC monolayer (Supplementary Fig. 26). We found both Clusters 1 and 2 of the organoid endothelial population characterized nearly entirely as “endothelial cell of vascular tree” (84.9% and 75.3%, respectively) (Supplementary Fig. 26), whereas the iETV2-hiPSC monolayer populations of P1, P2, P3 only characterized as 31.0%, 61.7%, 84.1%, respectively (Supplementary Fig. 6).

Furthermore, we found the endothelial cells to closely interact with the epithelial cells of the kidney organoid (Fig. 2d, 3b). Moreso, on scanning electron microscopy, we identified that the endothelial cells that encased the podocytes display a fenestrated cell membrane (Fig. 6h). Additionally, endothelial cells of the vascularized kidney organoid contain high expression of *PLVAP,* marker of fenestrated endothelial cells associated with bridging diaphragms of fenestrae and caveolae^67^ (Fig 6i). Fenestrations play a critical role in glomerular filtration and thus, this represents a significant novel finding in the kidney organoid^68^. It has been shown that ETV2 can play a role in functionally ‘resetting’ endothelial cells that adjusts and conforms to organoids in a tissue-specific manner^13^, and thus, this data demonstrates that iETV2-hiPSCs undergo further maturation and organ-specific differentiation within a kidney organoid niche. Taken together, our data demonstrates that the iETV2-hiPSCs are receptive to the microenvironmental cues of developing kidney organoids. As such, the resulting endothelial network yields a kidney-specific endothelial vascular tree alongside more mature epithelial cells.

## Discussion

Endothelial niches play a central role during organogenesis, controlling fate maturation, patterning and morphogenetics events^1–6^. However, iPSC-derived organoids grown *in vitro* frequently lack a well-developed endothelial niche which, in addition to promoting tissue maturation, is likely to be needed in future transplant settings where organ-specific endothelial may be functionally important. The failure to identify a common culture media that can support the development of both resident parenchymal epithelia and endothelial populations has complicated this effort. Another approach has been the addition of endothelial-promoting cytokines such as VEGF-A to the organoid differentiation medium, but while this expands angioblast cell numbers, it fails to support the formation of an integrated vascular network^20^. Strategies outside of media supplementation include the use of fluidic culture systems that promote vascular growth and survival^19–21^, and transplantation into immunodeficient mice, where the host vessels invade the xenograft^24,25^. However, these are cumbersome, low throughput and technically complicated to set-up for routine organoid experiments.

Our strategy took advantage of ETV2, a ‘pioneer’ transcription factor that instructs endothelial fate^13,14,69,70^, to genetically engineer in a doxycycline-inducible fashion, an endothelial niche. We achieved this by mixing an optimal ratio of iETV2-hiPSC to non-transgenic hiPSCs together, thus allowing endothelial cell numbers to be ‘tuned’ to a physiologically relevant quantity, while not overwhelming the organoid with vasculature. The resulting engineered niche is orthogonal to the organ-specific developmental program, but operates in tandem, offering an innovative, ‘plug-and-play’ approach to vascularize organoids.

In our study, we show for the first time, that endothelial cells generated from iETV2-hiPSCs develop a fenestrated, endothelial network capable of morphologically integrating with tubular cells, podocytes, and interstitial cells. This vascularization leads to a more mature podocyte population with interdigitated foot processes, endothelial glomerular invasion, and basement membrane maturation. The vascularized kidney organoids also developed a novel, integrated, and responsive renin cell population in the interstitium. Finally, we demonstrated that the development and maturation of these epithelial cells also coincides with the formation of a kidney-specific endothelial population, demonstrating the potential co-development and crosstalk occurring between the epithelial, interstitial, and endothelial cells in generating a heterogenous mature organoid.

This study represents the first utilization of iETV2-hiPSCs line to vascularize human kidney organoids. The protocol of iETV2-hiPSC:wildtype hiPSC was selected as to allow for a significant number of endothelial cells to incorporate into the kidney organoid in a physiologically relevant quantity, while not overwhelming the culture with endothelial cells. It is possible that with greater tuning of the ratio of cells, the doxycycline concentration, the doxycycline length of exposure and timing, different compositions of cell types within the kidney organoid can be achieved. We observed by snRNAseq there was a slightly diminished proportion of tubular cells in the vascularized kidney organoid, particularly distal tubule cells. Despite this observation, we found in qPCR and snRNAseq elevated expression levels of *GATA3*, a common distal tubule marker in kidney organoids. GATA3, however, has been shown to play a role in mesangial and podocyte development^71–73^. This may indicate that despite a decrease in number of distal tubule cells, there is an overall enhancement in the interstitial and podocyte populations. With greater tuning of protocol parameters as has been demonstrated in other studies, it may be possible to proximalize or distalize the vascularized organoids^74^. Finally, we observed the INT-2 interstitial population predominantly originates from the vascularized kidney organoid. Like other organoid studies these cells have been found to contain neural markers such as *DCX*, *NEUROD4*, and *STMN2* and typically are written off as unassigned or unidentifiable. Without better understanding of the heterogeneity observed in human kidney interstitia, it is still unknown the exact function or reason for this observation. Future characterization of this cell type with further immunostaining and functional studies would greatly benefit the understanding of the interstitial component of the kidney organoid.

Endothelial cells are also known to signal to surrounding tissues^75^. In the case of the developing glomerulus, disruption of the glomerular endothelium prevents podocyte maturation with a reduction in foot process and slit diaphragms^76^. In support of this, our vascularized organoids display glomeruli-like structures, comprised of juxtaposed podocytes and endothelial cells, with the podocytes forming foot processes and interdigitations, all of which are rare or absent in control organoids. The podocytes in vascularized organoids also express higher levels of mature podocyte markers^47,48^ such as *PTPRO*, *NPHS2*, *COL1A2*, *COL4A5, LAMC1, LAMA5 and CCN2,* compared to the unvascularized controls. Similar to other reports^47^, we found higher expression of some early podocyte markers in our vascularized organoids, compared to the controls, indicating that optimization of the culture conditions and/or state of endothelial maturation is needed to further improve endothelial-podocyte crosstalk. These markers have even persisted following long term transplantation^47^. To further mature kidney organoid podocytes, it may be necessary to direct the differentiation of ETV2-induced endothelial cells into the distinct glomerular endothelial cell type. This could be achieved with the introduction of fluid flow or alternatively, the identity of glomerular endothelial cells could be genetically induced, similar to our iETV2 strategy, by the inclusion of the transcription factors GATA5 and TBX3 that drive this fate^77^.

In investigating the emergence of the renin+ cell population in the interstitial of the vascularized kidney organoid, there are still significant questions that remain. The exact composition of the juxtaglomerular apparatus and whether the renin cells interact with other mesangial cells to juxtapose the podocytes and endothelial cells in the kidney organoid to form such an apparatus is unknown. We observe that with forskolin stimulation there is an elevated expression level of *REN,* however, in future studies we aim to investigate whether renin is successfully secreted from these cells, and whether these cells are functionally responsive to fluid pressure as occurs *in vivo*.

Work remains to fully understand the endothelial cells within the vascularized kidney organoid. During development, each tissue develops a vascular network with different types of vessels including arteries, veins and capillaries that are functionally specialized to each particular organ^67^. In our vascularized organoids we found little evidence that the endothelial cells underwent arteriovenous differentiation. This is most likely because the organoids were grown under static conditions, whereas *in vivo* the developing vasculature is highly responsive to blood flow, which helps remodel the immature plexus and induce different arteriovenous fates^78,79^. The addition of flow or shear stress to our current organoid protocol may be sufficient to achieve a more mature vascular network. Despite this, the endothelial cells adopted a transcriptional signature that is most similar to that of endogenous endothelia from the adult kidney. This suggests that the organoid microenvironment such as epithelial and interstitial cells are providing differentiation signals to the vascular plexus. As the nature of these signals has not been well characterized, our vascularized organoids offer a new tool to investigate these processes. This vasculogenic iPSC line can easily be adapted by existing fluidic protocols to enable perfusion, which may further mature the endothelial network and surrounding parenchymal cells^19,20^.

Overall, the kidney organoids generated with the use of the inducible endothelial niche contain a vascular network that morphologically integrates with podocytes, interstitial cells, and renin cells in a way that will allow investigation of disease processes that are often determined on a histological basis. Therefore, the work within this study moves the field of kidney tissue engineering a significant step forward to more adequately model disease of glomerular basement formation, disease of the juxtaglomerular apparatus, and with additional engineering modules – modelling of physiological conditions such as blood pressure or cyst formation. This inducible endothelial progenitor iPSC line was easily integrated into an established kidney organoid protocol, and thus demonstrates the potential for other organ systems and models to include an inducible vascular niche^3,6,13,14,80–82^. Our study lays groundwork for an easy to use and widely applicable engineering method for manipulation and augmenting a tissue vascular niche during *in vitro* human kidney organogenesis.

## Online Methods

### Generating ETV2-Inducible hiPSCs

rtTA expressing PGP1 hiPSCs previously generated^83^ were transfected, as we and other groups have done^28,84,85^, using Lipofectamine 3000 (Thermo Fisher Scientific, L3000001) with Super PiggyBac Transposase (System Biosciences, PB210PA-1) and the PiggyBac transposon vector with hETV2-2A-EGFP under control of the tetracycline responsive element promoter. Transfected cells were selected by adding 0.5mg/mL puromycin to mTeSR1 maintenance medium (STEMCELL Technologies, 85850). PGP1-ETV2 hiPSCs were karyotyped using Thermo Fischer Scientific Karyostat+ Karyotyping service.

#### Generating HNF4A-GATA3-MAFB Triple hiPSCs

HNF4A^mCitrine^:MAFB^mTagBFP2^:GATA3^mCherry^ (Triple) reporter iPSCs were generated by incorporating the mCitrine gene at the start codon of the HNF4A locus in the previously described dual reporter iPSC line, MAFB^mTagBFP2^:GATA3^mCherry86^. In vitro transcribed mRNA encoding the SpCas9-Gem variant^87^, plasmid encoding a sgRNA targeting the 5’ end of the HNF4A locus and a gene targeting plasmid encoding a mCitrine-T2A gene cassette flanked by 485 bp and 708 bp homology arms corresponding to sequences directly upstream and downstream of the HNF4A locus respectively, were introduced into iPSCs using the Neon Transfection System as previously described^86^ (Supplementary Table 4). To identify correctly targeted iPSCs, genomic DNA was isolated using the DNeasy Blood & Tissue Kit (QIAGEN) and PCR analysis was performed using primers that flank the 5′ and 3′ recombination junction. A clone containing homozygous insertion of the mCitrine reporter, determined by PCR using primers that flank the intended target site, was selected for further expansion and characterization.

### Cell Culture

Kidney organoids were generated using the previously published MANZ2-2 hiPSC line^88^ and the HNF4A-GATA3-MAFB (Triple) hiPSC line. These cells were combined with PGP1-ETV2 hiPSCs to generate vascularized human kidney organoids. All hiPSCs were cultured between 10 and 60 passages. MANZ2-2 and Triple hiPSCs were cultured in mTeSR1 medium while PGP1-iETV2-hiPSCs were cultured in NutriStem XF/FF culture medium (Reprocell, 010005). All hiPSCs were cultured in media supplemented with 1% Penicillin-Streptomycin (Pen-Strep) (Gibco, 15140122) and 0.01% Plasmocin (InvivoGen, ant-mpp-1). hiPSCs were passaged and cultured on tissue-culture plates coated with Cultrex basement membrane extract (R&D Systems, 343401002). For passaging, hiPSCs at 70-80% confluency were washed in DPBS without calcium or magnesium (Gibco, 14190144), then incubated at 37°C in 2mL of gentle cell dissociation reagent (GDR) (STEMCELL technologies, 100-0485). GDR was aspirated and cells were lifted from tissue-culture plates using cell lifters (Fisher Scientific, 08-100-240). Cells were then resuspended in media and diluted into new tissue culture plates at a ratio of 1:3-1:8. Cells were then maintained in mTeSR1 or NutriStem with daily media changes. For induction of endothelial cascade with PGP1-iETV2-hiPSCs, NutriStem was supplemented with 0.5μg/mL doxycycline (hyclate) (STEMCELL technologies, 72742) unless stated otherwise in the Fig. s.

### Kidney Organoid Generation

Kidney organoids were generated using a previously published protocol^30,31^. In summary, hiPSCs were washed twice in DPBS, then colonies were dissociated with GDR, and resuspended in Stage I media which is comprised of TeSR-E5 media (STEMCELL Technologies, 05916) supplemented with 0.1% (v/v) Insulin Transferrin Selenium Ethanolamine (Gibco, 51500056), 1% (v/v) Pen-Strep, 0.25% (v/v) poly(vinyl alcohol) (PVA) (Millipore Sigma, P8136), 0.01% (v/v) Plasmocin, 8μM CHIR99021 (STEMCELL Technologies, 72054), 3.3μM Y27632 (STEMCELL Technologies, 72304), and 0.1mM beta-mercaptoethanol (Fisher Scientific, 21-985-023). Cell colonies in suspension were transferred into ultra-low attachment 6 well plates (Corning, CLS3471-24EA) and cultured for 2 days on a gyrating rocking platform. On day 2, half of the media was aspirated and replaced with the same TeSR-E5 media with supplements omitting beta-mercaptoethanol and Y27632 and cultured for 24hrs to form embryoid bodies. On day 3, embryoid body spheroids were filtered by size with a 200μm PluriStrainer (Fisher Scientific, NC1474108), washed in low-glucose DMEM (Gibco, 11054001), and transferred to Stage II media which is comprised of low-glucose DMEM supplemented with 10% (v/v) KnockOut™ Serum Replacement (KOSR) (Gibco, 10828028), 1% (v/v) Penn-Strep, 1% (v/v) non-essential amino acids (NEAA) (Gibco, 11140050), 1% (v/v) HEPES (Gibco, 15630080), 1% (v/v) GlutaMAX™ (Gibco, 35050061), 0.01% (v/v) Plasmocin, and 0.25% (v/v) PVA. Organoids were then continually grown on a gyrating rocker with media changes every other day until day 18. To generate vascularized kidney organoids, the PGP1-ETV2 hiPSC line was combined with a control hiPSC line (MANZ2-2 or Triple) at a ratio of 1:5 on day 0 in Stage I media. The same protocol for organoid formation was carried out with the addition of 0.5μg/mL of doxycycline to Stage II media from day 5-18, supplemented each day, unless otherwise stated.

### Whole Mount Clearing and Immunofluorescence

Organoids were fixed in 4% paraformaldehydes (PFA) (Fisher Scientific, AC416785000) for 24hrs and rinsed in PBS three times. Organoids were then rendered optically clear through an adapted version of the “Clear, Unobstructed Body Imaging Cocktails and Computational-analysis” (CUBIC) clearing protocol^89–91^. In summary, organoids were first placed into 1:1 dH_2_O:CUBIC R1, with three times replacement of liquid every 2 hours followed by a 24hr incubation at room temperature. CUBIC R1 is comprised of 35% (wt/v) dH_2_O, 15% (wt/v)Triton X-100 (Sigma-Aldrich, T8787), 25% (wt/v)N,N,N’,N’-Tetrakis(2-Hydroxypropyl)ethylenediamine (quadrol) (Sigma-Aldrich, 122262), and 25% (wt/v)urea (Thermo Scientific, 29700). Organoids were then washed three times in IHC Buffer which was comprised of 500mL PBS, 0.1% (v/v) Triton X-100, 0.5% (v/v) bovine serum albumin (BSA) (Tocris Bioscience, 5217), and 0.01% (v/v) Sodium Azide (Sigma-Aldrich, 71289). Organoids were then washed a final time and incubated over night at room temperature. Next, organoids were incubated at room temperature for 48 hours in IHC Buffer supplemented with primary antibody (Supplementary Table 1). Organoids were then washed three times in IHC Buffer, and incubated at room temperature over night in IHC Buffer supplemented with secondary antibody (Supplementary Table 2). Organoids were then washed three times in PBS with a final PBS wash overnight. A 0.2% (wt/v) agarose solution was then made by dissolving 0.4g of low-melting point agarose (Invitrogen, 16520100) in 10mL of dH_2_O at 55°C. 10mL of CUBIC R2 which was comprised of 15% (wt/v) dH_2_O, 0.1% (v/v) Triton X-100, 25% (wt/v) urea, and 50% (wt/v) sucrose (Fisher Scientific, AA36508A1) was then slowly added. Organoids were then thoroughly removed of PBS and resuspended in the 0.2% molten agarose-CUBIC R2 solution. The molten solution with organoids suspended within, was then transferred to a 200μL Combitips® tube (Eppendorf, 0030089774) with the tip removed. The molten gel-organoid-suspension was transferred to 4°C for 15 minutes to allow for solidification. Using the plunger of the Combitips®, the solid tube of agarose with organoids embedded within, was then pushed out into a 24-well glass bottom plate (Fisher Scientific, NC0397150). ∼5μL of ultra-adhesive glue was then placed at both ends of the agarose tube to secure the tube to the bottom of the well. The glue was allowed to dry for 5 minutes, and the well was then filled with CUBIC R2. The liquid was replaced every hour, 3 times, and left to incubate at room temperature overnight. Organoids were then imaged on a Nikon A1R Spectral Confocal microscope. Three-dimensional renderings of the organoids were then generated using Bitplane Imaris v10.0.0 (Oxford Instruments).

### Angiotool analysis of vascular networks

Maximum intensity z-stack projections of the PECAM1 channel of hiPSC monolayer or 300μm depths of organoids were generated. The z-stack projections were converted to a maximum intensity composite image in FIJI (ImageJ). Default settings were kept in AngioTool^39^, and a vessel diameter of 4, 7, 10, 14μm was used for analysis of all samples as has been done in other published studies^19^. Quantification of vascular networks were analyzed and graphed in GraphPad Prism. A non-parametric t-test (Mann-Whitney) was employed to analyze statistical significance between control and vascularized conditions.

#### Immunofluorescence quantification

Individual cells in immunofluorescence were quantified for their per cell immunofluorescent intensity through an adapted protocol of previously published work^92^. In summary, immunofluorescent images of organoids with DAPI, EGFP, MCAM and PECAM1 were acquired and analyzed in FIJI. Background fluorescence was adjusted in all channels and images by using Otsu Dark threshold with minimum threshold set to 600 pixel intensity units. This nuclear region was then subtracted from EGFP, MCAM and PECAM1 images to obtain nuclei-free fluorescent images. The total pixel intensity of the non-nuclear regions was calculated for each channel, then divided by the nuclei count to obtain a final value of average summed pixel intensity per cell. Quantification of per cell immunofluorescent intensity was analyzed and graphed in GraphPad Prism. A non-parametric ANOVA test with multiple comparisons (Kruskal-Wallis) was employed to analyze statistical significance between protein immunofluorescence at various timepoints.

#### Electron Microscopy

Organoids were harvested at day 18 for scanning and transmission electron microscopy (SEM/TEM). Organoids were placed in 2.5% glutaraldehyde fixative for 60 minutes for morphological preservation. For SEM, organoids were washed in PBS, and submerged in 1% osmium tetroxide for 60 minutes. The organoids were then dehydrated through a series of 30-100% ethanol washes, then washed with hexamethyldisalazane (HDMS) and allowed to dry. Then, organoids were placed onto aluminum stubs with conductive copper double sided adhesive tape, sputter coated with gold/palladium and placed in a JSM-6335F field emission scanning electron microscope at 3 kV (JEOL, Tokyo, Japan). For TEM, glutaraldehyde fixed organoids were washed in PBS, and submerged in 1% osmium tetroxide and 1% potassium ferricyanide for 60 minutes. The organoids were then dehydrated in a series of 30-100% ethanol washes then embedded in Polybed 812 resin. Once the resin was cured at 37o overnight then at 65o for 2 days. Sixty nm sections were sectioned using a diamond knife and placed onto 200 mesh copper grids. Sections were imaged using a JEOL JEM 1400 Flash transmission electron microscope (Peabody, MA) at 80 kV, and photographed with a bottom-mount AMT 2k digital camera (Advanced Microscopy Techniques, Danvers, MA).

#### RNA extraction and RT-qPCR

RNA was extracted from monolayers of hiPSCs or whole kidney organoids using TRI Reagent™ (Thermo Fisher Scientific, AM9738) at room temperature for 30 minutes with intermittent agitation. Samples were then either stored at -80°C for storage, or immediately processed using PureLink™ RNA Mini Kit (Invitrogen, 12183018A). RNA quality was analyzed using NanoDrop 1000 (Thermo Fisher Scientific) for purity and concentration. RNA was stored at -80°C for long-term storage, or immediately processed using qScript™ cDNA Supermix (Andwin Scientific, 101414106). cDNA was stored at -80°C for long-term storage. RT-qPCR experiments were performed in 96-well plates (Applied Biosystems, 4346906) using SYBR™ Green Master Mix (Applied Biosystems, A25778) forward and reverse primers were generated by Integrated DNA Technologies (IDT) (Supplementary Table 3). RT-qPCR was processed on QuantStudio 12K Flex PCR System (Thermo Fisher Scientific). Ct values were processed according to previously published protocols^93^ and graphs were generated in Prism (GraphPad, 9.1.0).

### Single-cell RNA sequencing of ETV2-hiPSC line

iETV2-hiPSCs were prepared as described by the 10x Genomics Single Cell 3 ′ v2 Reagent Kit user guide. After 4 days of doxycycline exposure, a monolayer of cells were incubated with trypsin for 10 minutes at 37 ° C, followed by gentle pipetting using a serological pipette to dislodge and dissociate aggregates. Samples were washed in PBS lacking calcium and magnesium ions (−/−) + 0.04% BSA twice and re-suspended at a final concentration of 1000 cells/μL in PBS −/− + 0.04% BSA. Using Trypan Blue, a live cell count was performed to identify dead cells. Following cell counting, the cells and 10x Genomics reagents were loaded into the single cell cassette, with a target of 7500 cells single cells for analysis, accounting for predicted cell loss and doublets as laid out in the 10X Genomics Chromium Single Cell 3’ Reagent Kits User Guide (v3.1 Chemistry Dual Index) with Feature Barcoding technology for Cell Multiplexing, User Guide, CG000388. After generation of GEMs, the cDNA library was prepared by University of Pittsburgh Single Cell Core staff following the appropriate steps determined by the 10x Genomics user guide. Libraries were sent to UPMC Genome Center for sequencing on a NovaSeq 6000. The 10x Genomics CellRanger pipeline was used to align reads to the reference genome (Hg38) appended with transgene sequences, to assign reads to individual cells, and to estimate gene expression based on UMI counts^94^. Utilizing the Seurat package^95^, cells were filtered out if they contained less than 200 unique molecular identifiers, contained less than 200 genes, and had more than 5% counts from mitochondrial genes. Counts were then log-normalized using Seurat normalization function. Mutual nearest neighbors and clusters were then generated using Seurat. Downstream analysis and graphs were then generated with Seurat “DimPlot”, “VlnPlot”, “DotPlot” and “FeaturePlot”. Differential gene expression between clusters was calculated using the ‘vst’ method in Seurat. Gene ontology analysis was then performed using Enrichr^96,97^, and Gene Ontology (GO)^98,99^. Single cell data for Tabula Sapiens endothelial cells^35^ was acquired from CZ CELLxGENE data portal. Endothelial cells from this dataset were parsed by organ of origin and gene ontology term. This dataset was then used as a training set in SingleCellNet^36^ and aligned to the PGP1-iETV2-hiPSC scRNAseq dataset as a query. Pseudotime trajectory analysis was performed using Monocle3^100^ by setting Cluster 0 as a starting point and utilizing ETV2 and EGFP as trajectory genes.

#### Single-nuclear sequencing of kidney organoids

Day 18 MANZ2-2 control and vascularized kidney organoids were washed in DPBS once, then transferred into 37°C 0.25% trypsin-EDTA (Gibco, 15400054) and incubated with intermittent agitation for 10 minutes to obtain single cells. Cells were then centrifuged at 800rpm for 5 minutes and resuspended in DPBS and filtered through a 40μm cell filter (pluriSelect, 43-50040-51). 10X Genomics CG000365 Demonstrated Protocol was then adapted to isolate nuclei. In summary, cells were treated with chilled Multiome Lysis Buffer on ice for 5 minutes to obtain nuclei. Nuclei were centrifuged at 500 rcf for 5 minutes, supernatant was removed and resuspended in Wash Buffer. Nuclei were washed and centrifuged 3 additional times. Finally, supernatant was removed, and nuclei were resuspended in Nuclei Buffer. Composition of all buffers can be found in Supplementary Tables 5, 6 and 7.

Using Trypan Blue, a live cell count was performed to identify dead cells. Following counting, cells and 10x Genomics reagents were loaded into the single cell cassette, with a target of 1000 nuclei for analysis, accounting for predicted cell loss and doublets as laid out in 10X Genomics Chromium Single Cell 3’ Reagent Kits User Guide (v3.1 Chemistry Dual Index), User Guide, CG000315. After generation of GEMs, the cDNA library was prepared by University of Pittsburgh Single Cell Core staff following the appropriate steps determined by the 10x Genomics user guide. Libraries were sent to UPMC Genome Center for sequencing on a NovaSeq 6000. Raw reads were processed using Partek® Flow® software, v10.0 and aligned to the GRCh38 human reference genome. The same downstream computational analysis pipeline from the previous methods section on scRNAseq was then carried out on this snRNAseq population with some additional metrics. To account for batch variation in samples, the Harmony package was deployed to synergize control and vascularized datasets^101^. Cellular identities were confirmed using DevKidCC^41^. In addition to the Tabula Sapiens comparison using SingleCellNet, the PGP1-ETV1 hiPSC scRNAseq dataset was also used as a training set with SingleCellNet to query the endothelial cells of the control and vascularized kidney organoid.

#### Renal Capsule Implantations

Human kidney organoids were implanted under mice renal capsule using a procedure adapted from previously published methods^24,102^. Male NSG mice (Jackson laboratories) at 8-10 weeks old were anesthetized with isoflurane. The animals were immobilized in a dorsal lateral position, and under continuous anesthesia influx, the fur was shaved, and the skin was cleansed and disinfected with ethanol wipes and iodine. A skin incision was performed in the lower left flank followed by a mucosa incision upon localization of the left kidney. After externalizing the kidney, saline was used to keep it moist, and gentamicin was applied locally with a cotton swab to prevent infection. An adapted 27G scalp was aseptically cut at a 45 degrees angle. The extremity closer to the needle was used to create a nip on the kidney capsule, forming the letter L. The other portion was attached to a 1 mL syringe, which collected vascularized organoids from a petri dish. The organoids were pushed to the other extremity and injected underneath the kidney capsule. The area was then cauterized, the kidney was delicately pushed back into the abdominal cavity, the mucosa aseptically sutured and the skin stapled. The same procedure was performed on the lower right flank of each animal, but non-vascularized organoids were injected instead. Animals were returned to their cages after the procedure, water and food were provided *ad libitum*, and euthanasia occurred 28 days post-implantation. Kidneys were collected for histological analysis.

### Data Availability

The data supporting the findings of this study are openly available in GEO under accession number GSE232767.

---

**Supplementary Table 1:** Primary immunostaining antibodies.

| ***Name*** | ***Host Animal*** | ***Vendor*** | ***Catalog Number*** | ***Dilution*** |
| --- | --- | --- | --- | --- |
| MCAM | Rabbit | Abcam | ab75769 | 1:200 |
| PECAM1 | Mouse | Abcam | ab9498 | 1:200 |
| HNF4A | Rabbit | Abcam | ab92378 | 1:200 |
| CDH1 | Rabbit | Abcam | ab40772 | 1:200 |
| NPHS1 | Guinea Pig | Progen | GP-N2 | 1:200 |
| REN | Rabbit | Abcam | ab212197 | 1:200 |
| NRP1 | Goat | R&D Systems | AF566 | 1:200 |
| MEIS1/2/3 | Mouse | Fisher Sci. | 50-199-2860 | 1:200 |
| EMCN | Rat | Abcam | ab106100 | 1:200 |

**Supplementary Table 2:** Secondary immunostaining antibodies.

| ***Target Animal*** | ***Host Animal*** | ***Fluorophore*** | ***Vendor*** | ***Catalog Number*** | ***Dilution*** |
| --- | --- | --- | --- | --- | --- |
| Mouse | Donkey | AF647 | Thermo Fisher | A-31571 | 1:250 |
| Rabbit | Donkey | AF594 | Thermo Fisher | R37119 | 1:250 |
| Guinea Pig | Goat | DyLight 594 | Thermo Scientific | SA510096 | 1:250 |
| Goat | Rabbit | AF594 | Abcam | ab150144 | 1:250 |
| Rat | Goat | AF594 | Invitrogen | A-11007 | 1:250 |
| -- | -- | DAPI | Invitrogen | D1306 | 1:250 |

**Supplementary Table 3:**
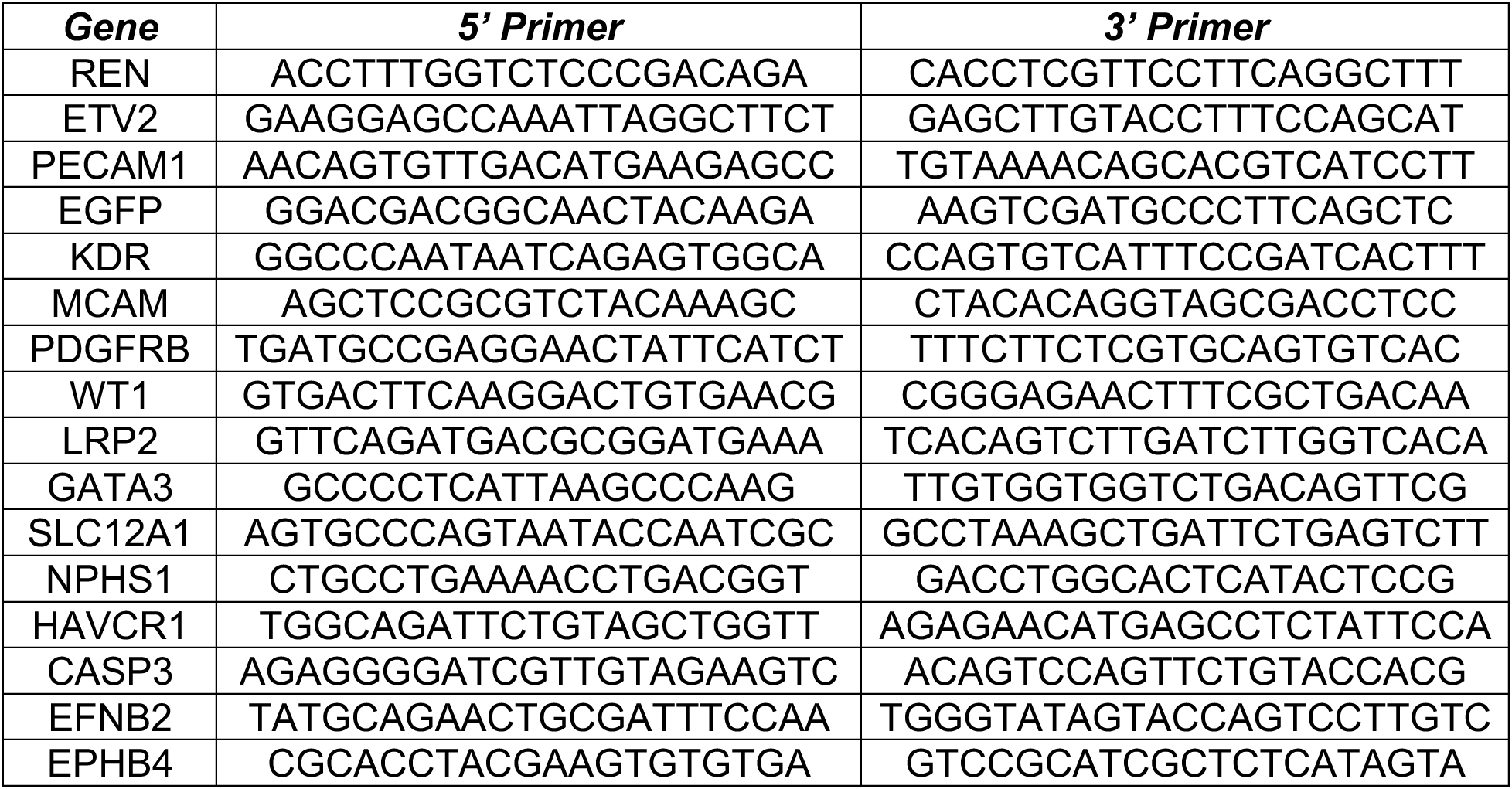
Primers for RT-qPCR.

**Supplementary Table 4:**
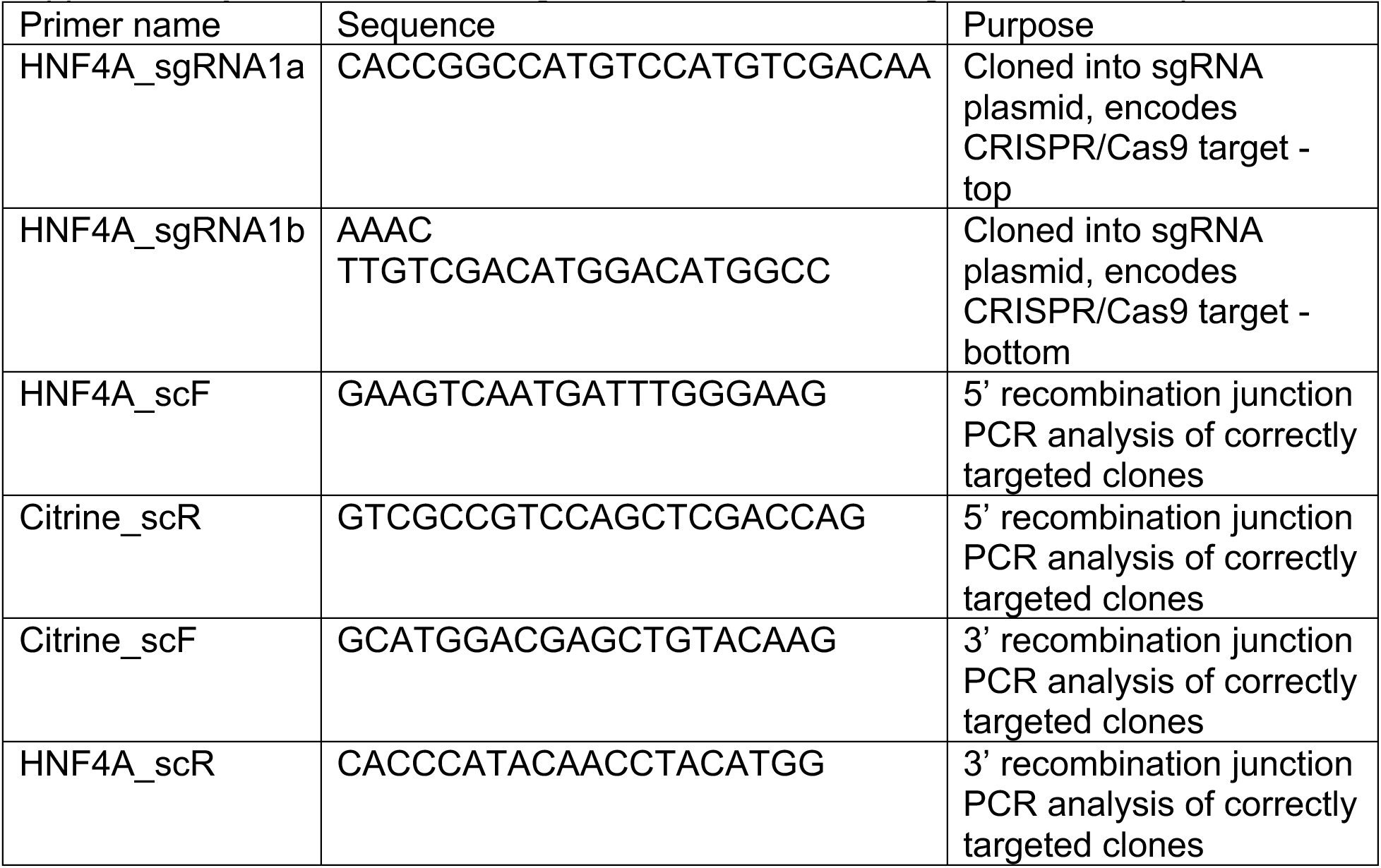
List of Oligodendronucleotides for generation of Triple

**Supplementary Table 5:** Multiome Lysis Buffer Composition.

| ***Reagent*** | ***Stock*** | ***Final*** | ***Amount*** |
| --- | --- | --- | --- |
| Tris-HCl (pH 7.4) | 1M | 10 mM | 20 µl |
| NaCl | 5M | 10 mM | 4 µl |
| MgCl<sub>2</sub> | 1M | 3 mM | 6 µl |
| Tween-20 | 10% | 0.1% | 20 µl |
| Nonidet P40 Substitute | 10% | 0.1% | 20 µl |
| Digitonin | 5% | 0.01% | 4 µl |
| BSA | 10% | 1% | 200 µl |
| DTT | 1000 mM | 1 mM | 2 µl |
| RNase inhibitor<br>40 U/µl | 40 U/µl | 1 U/µl | 50 µl |
| Nuclease-free<br>water | - | - | 1.67 mL |

**Supplementary Table 6:** Wash Buffer Composition.

| ***Reagent*** | ***Stock*** | ***Final*** | ***Amount*** |
| --- | --- | --- | --- |
| Tris-HCl (pH 7.4) | 1 M | 10 mM | 40 µl |
| NaCl | 5 M | 10 mM | 8 µl |
| MgCl<sub>2</sub> | 1 M | 3 mM | 12 µl |
| BSA | 10% | 1% | 400 µl |
| Tween-20 | 10% | 1% | 40 µl |
| DTT | 1000 mM | 1 mM | 4 µl |
| RNase inhibitor | 40 U/µl | 1 U/µl | 100 µl |
| Nuclease-free<br>Water | - | - | 3.40 mL |

**Supplementary Table 7:** Diluted Nuclei Buffer Composition.

| ***Reagent*** | ***Stock*** | ***Final*** | ***Amount*** |
| --- | --- | --- | --- |
| Nuclei Buffer<br>(20X) | 20X | 1X | 50 µl |
| DTT | 1000 mM | 1 mM | 1 µl |
| RNase inhibitor | 40 U/µl | 1 U/µl | 25 µl |
| Nuclease-free<br>Water | - | - | 924 µl |

**Supplementary Fig. 1: Normal karyotype for engineered doxycyline inducible iPSC line. (a)** Whole genome view displaying all somatic and sex chromosomes in one frame. A value of 2 represents a normal copy number state. A value of 3 represents chromosomal gain. A value of 1 represents a chromosomal loss. The pink, green, and yellow colors indicate the raw signal for each individual chromosome probe, while the blue represents the normalized probe signal which is used to identify copy number and aberrations, if any.

**Supplementary Fig. 2: Doxycycline response of hiETV2-iPSC endothelial differentiation.** Representative qPCR results of ETV2 iPSC induced cell line after 4 days of exposure, ETV2 and PECAM1. Error bars represent standard deviation of two technical replicates.

**Supplementary Fig. 3: Graphical schematic of per cell fluorescence intensity quantification using ImageJ FIJI.**

**Supplementary Fig. 4: Immunofluorescence of iETV2-hiPSCs.** Representative imaging of DAPI, GFP, ERG, PECAM1, iETV2-hiPSCs post-exposure to four days of doxycycline.

**Supplementary Fig. 5. scRNAseq featureplots of iETV2-hiPSCs.** Featureplots of differentially expressed genes for P1, P2, P3 clusters.

**Supplementary Fig. 6: SingleCellNet Characterization of iETV2-hiPSC scRNAseq. (a)** iETV2-hiPSC classification using Tabula Sapiens organ-specific endothelial classifiers. **(b)** iETV2-hiPSC classification using Tabula Sapiens endothelial gene ontology classifiers.

**Supplementary Fig. 7: SingleCellNet using Endothelial Tabula Sapiens by organ system. (a)** Receiver operating characteristic (ROC) analysis demonstrating classification power of Tabula Sapiens Endothelial Dataset by organ system. **(b)** Classification of cell types by cell type classifier. **(c)** Heatmap classification of cell types with cell type classifier. **(d)** Top gene pairs determined by SingleCellNet for Tabula Sapiens endothelial organ specific dataset.

**Supplementary Fig. 8: SingleCellNet using Endothelial Tabula Sapiens Gene Ontology Terms. (a)** ROC analysis demonstrating classification power of Tabula Sapiens Endothelial Dataset classifiers with gene ontology terms. **(b)** Heatmap classification of classifiers. **(c)** iETV2-hiPSC scRNAseq data queried in SingleCellNet to Tabula Sapiens Endothelial gene ontology classifiers. **(d)** Top gene pairs determined by SingleCellNet for Tabula Sapiens Endothelial gene ontology classifiers.

**Supplementary Fig. 9: Arterial and Venous markers of differentiated ETV2 iPSCs. (a)** EFNB2, EPHB4 and combined expression based on UMAP. **(b)** EFNB2 expression plotted against EPHB4 expression (r = 0.07).

**Supplementary Fig. 10: Optimization of doxycycline for vascularization protocol. (a)** Altering doxycycline window starting point; representative qPCR results. **(b)** Altering doxycycline window length; representative qPCR results. **(c)** Altering doxycycline concentration; representative qPCR results. qPCR error bars represent standard deviation of two technical replicates.

**Supplementary Fig. 11: Optimization of iETV2-hiPSCs to wildtype iPSC ratio for vascularization protocol. (a)** Representative immunofluorescence images varying ratios of cellular composition. Scale bar = 200μm. **(b)** Representative qPCR. qPCR error bars represent standard deviation of two technical replicates.

**Supplementary Fig. 12: GFP / PECAM1 colabelling in kidney organoids with iETV2- iPSCs added. (a)** Representative quantification of GFP+ and PECAM1+ cells in kidney organoids. Scale bar = 200μm. **(b)** Representative images quantified for GFP+/PECAM1+ colabelling.

**Supplementary Fig. 13: Control and vascularized kidney organoid immunofluorescent images for endothelia and interstitia. (a)** Representative images of control and vascularized kidney organoids for PECAM1, endomucin, **(b)** NRP1; scale bar = 200μm. **(c)** Intersitial immunofluorescence with MEIS1/2/3.

**Supplementary Fig. 14: Triple hiPSC control and vascularized kidney organoids. (a)** Representative RT-qPCR of Triple hiPSC control and vascularized kidney organoids; error bars represent standard deviation of two technical replicates. **(b)** Representative immunofluorescent images of Triple control and vascularized kidney organoid with ETV2- GFP labelled endothelial cells and MAFB-BFP labelled podocytes.

**Supplementary Fig. 15: Control and vascularized kidney organoids implanted under renal capsule of NSG mice. (A)** Control human kidney organoids implanted under renal capsule with **(A’-A’’’)** representative insets. **(B)** Vascularized human kidney organoids implanted under renal capsule with **(B’-B’’’)** representative insets.

**Supplementary Fig. 16: DevKidCC classification of snRNAseq control and vascularized kidney organoid. (a)** LineageID Classification of snRNAseq organoid data. **(b)** DevKidCC classification of snRNAseq organoid data. **(c)** DevKidCC nephron classification scoring. **(d)** DevKidCC interstitia classification scoring. **(e)** DevKidCC LineageID scoring for control and vascularized kidney organoid by composition percentage. Labels: endothelial (ENDO), nephron progenitor cells (NPC), ureteric epithelia (UrEp), early nephron (EN), cortical stroma (CS), medullary stroma (MS), mesangial cells (MesS), ureteric outer stalk (UOS), ureteric inner stalk (UIS), early distal tubule (EDT), distal tubule (DT), Loop of Henle (LOH), early proximal tubule (EPT), parietal epithelial cell (PEC), early podocyte (EPod), podocyte (Pod).

**Supplemental Fig. 17: snRNAseq of control and vascularized tubular cells. (a)** UMAP of tubule cells from both control and vascularized kidney organoids. **(b)** DevKidCC Classification of tubular UMAP by proximal and distal tubule scoring. **(c)** UMAP colored by proximal and distal tubule according to DevKidCC. **(d)** UMAP of tubular cells classified by proximal and distal tubule subcomponents based on **(e)** top differentially expressed genes. **(f)** Proximal and distal tubule segments by canonical gene markers. **(g)** Definitive proximal tubule plotted on distinct UMAP with overlapping clusters and indistinct differentially expressed genes between control and vascularized kidney organoid. **(h)** Definitive distal tubule plotted on distinct UMAP with overlapping clusters and indistinct differentially expressed genes between control and vascularized kidney organoid. PPT: Proliferating Proximal Tubule; EPT: Early Proximal Tubule; PT: Proximal Tubule; DT: Distal Tubule; EDT: Early Distal Tubule.

**Supplementary Fig. 18: snRNAseq analysis of podocyte maturation.** VEGF signaling from podocytes. **(a)** Canonical genes of early and late podocyte maturation. **(b)** Canonical markers of glomerular basement membrane composition change during maturation. **(c)** CellChat quantification of VEGF signaling in control and vascularized kidney organoid snRNAseq conditions. **(d)** FeaturePlots of signals part of VEGF pathway in podocyte snRNAseq clusters.

**Supplementary Fig. 19: Podocytes from vascularized kidney organoid predominant cluster gene set enrichment analysis.**

**Supplementary Fig. 20: Canonical genes of interstitial cell populations from interstitial control and vascularized kidney organoid snRNAseq.**

**Supplementary Fig. 21: Differential gene expression in clusters of interstitial cells on snRNAseq.** For condition: red = vascularized, blue = control.

**Supplementary Fig. 22: *REN* and *EGFP* expression in control and vascularized snRNAseq interstitia.**

**Supplementary Fig. 23: Light microscope images of forskolin stimulated control and vascularized kidney organoids.** White arrows indicate cyst formation in the vascularized kidney organoid with forskolin exposure. Scale bar = 100μm.

**Supplementary Fig. 24: SingleCellNet training with iETV2-hiPSC scRNAseq dataset. (a)** ROC analysis of clusters 0 (P1), 1 (P2), 2 (P3). **(b)** Heatmap and **(c)** classification plots of classifiers. **(d)** Top gene pairs determined by SingleCellNet for classification. **(e)** Endothelial cells of kidney organoids snRNAseq queried to iETV2-hiPSC endothelial like cells using SingleCellNet.

**Supplementary Fig. 25: Ephrin localization in endothelial population of snRNAseq control and vascularized kidney organoid cells. (a)** FeaturePlots of EFNB2, EPHB4 and colocalization. **(b)** ENB2 vs EPHB4 plot of control and vascularized kidney organoid endothelial cells (r = 0.04).

**Supplementary Fig. 26: Morphological classification of endothelial cells in MANZ2-2 control and vascularized kidney organoid using SingleCellNet and Tabula Sapiens endothelial cell gene ontology classifier.**

## Acknowledgements

N.A.H. discloses support for the research of this work from the National Institute of Diabetes and Digestive and Kidney Diseases UC2 DK126122 and P30 DK079307, and by support from the Vascular Medicine Institute, the Hemophilia Center of Western Pennsylvania, and the Institute for Transfusion Medicine. M.R.E. discloses support for publication of this work by NIH center grant P30 DK120531 from the National Institute of Diabetes and Digestive and Kidney Diseases (to Pittsburgh Liver Research Center- PLRC); NIH grants R01 EB028532 from the National Institute of Biomedical Imaging and Bioengineering, R01 HL141805 from the National Heart Lung and Blood Institute and NSF grant (award number 2134999) (to M.R.E.). S.K discloses support for publication of this work by NIH center grant P30 DK120531 from the National Institute of Diabetes and Digestive and Kidney Diseases (to Pittsburgh Liver Research Center-PLRC); NIH grants R01 EB028532 and R01EB024562 from the National Institute of Biomedical Imaging and Bioengineering. J.C.M. discloses support for the research of this work, in part, from the University of Pittsburgh CATER training grant T32EB001026. A.W. discloses support for the imaging and instrumentation from 1S10OD019973-01. We acknowledge and thank Tracy Tabib, Heidi Monroe, and the team at the Single Cell Genomics Core of the University of Pittsburgh for performing scRNAseq and snRNAseq. A.P. would like to acknowledge and thank the Ben J. Lipps Research Fellowship Program for their support. C.C. and T.C. would like to acknowledge and thank the NIH (R01DK127634, RC2 DK125960) and CPRIT (RP220201) for supporting the research in this work.

