## Supplementary Figures for "Genetically engineering endothelial niche in human kidney organoids enables multilineage maturation, vascularization and de novo cell types"

*F: 412-383-2211; P: 412-648-9918*

1. Department of Developmental Biology, School of Medicine, University of Pittsburgh, Pittsburgh PA 15213, USA
2. Department of Pathology, Division of Experimental Pathology, School of Medicine, University of Pittsburgh, Pittsburgh PA 15213, USA
3. Pittsburgh Liver Research Center, University of Pittsburgh, Pittsburgh, PA 15261, USA
4. Department of Molecular Biology and Hamon Center for Regenerative Science and Medicine, University of Texas Southwestern Medical Center, Dallas, TX 75390, USA
5. Department of Internal Medicine, Division of Nephrology, University of Texas Southwestern Medical Center, Dallas, TX 75390, USA
6. Department of Pediatrics, School of Medicine, University of Pittsburgh, Pittsburgh PA, 15213
7. Center for Biologic Imaging, University of Pittsburgh, Pittsburgh, PA 15213, USA
8. Department of Bioengineering, Swanson School of Engineering, University of Pittsburgh, Pittsburgh, PA 15261, USA
9. Division of Nephrology, Department of Medicine, School of Medicine, Washington University in St. Louis, St. Louis, MO 63130
10. Murdoch Children's Research Institute, Melbourne, Victoria, Australia
11. Department of Anatomy and Neuroscience, The University of Melbourne, Melbourne, Victoria, Australia
12. Department of Paediatrics, The University of Melbourne, Melbourne, Victoria, Australia
13. Department of Developmental Biology, School of Medicine, Washington University in St. Louis, St. Louis, MO 63130
14. Department of Molecular Medicine and Pathology, University of Auckland, Auckland 1010, New Zealand
15. Department of Molecular Biology, UT Southwestern Medical Center, Dallas, TX 75390
16. McGowan Institute for Regenerative Medicine, University of Pittsburgh, Pittsburgh, PA 15219, USA

a

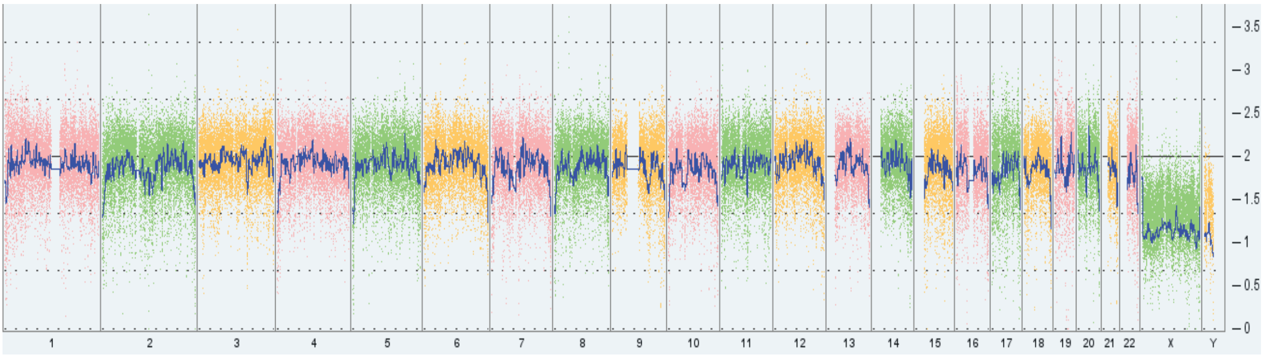


**Supplementary Fig. 1: Normal karyotype for engineered doxycyline inducible iPSC line.**

a

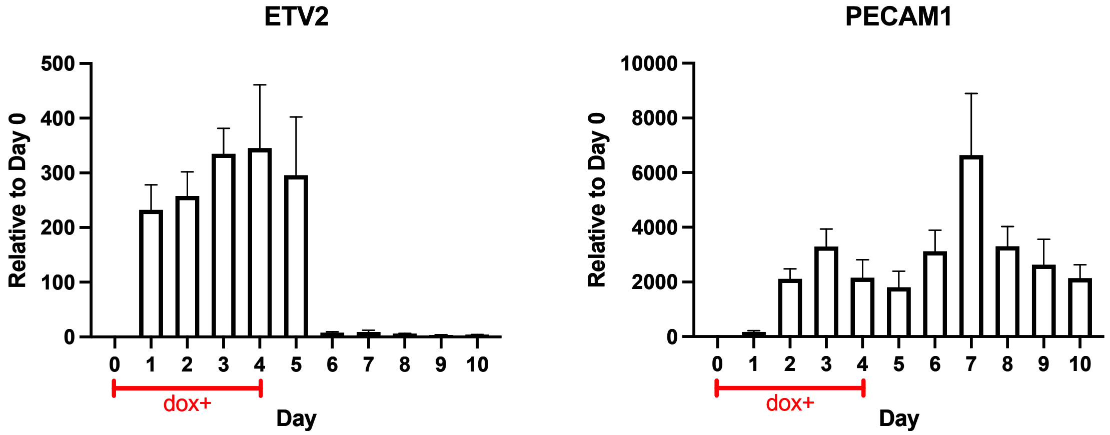


**Supplementary Fig. 2: Doxycycline response of hiETV2-iPSC endothelial differentiation.**

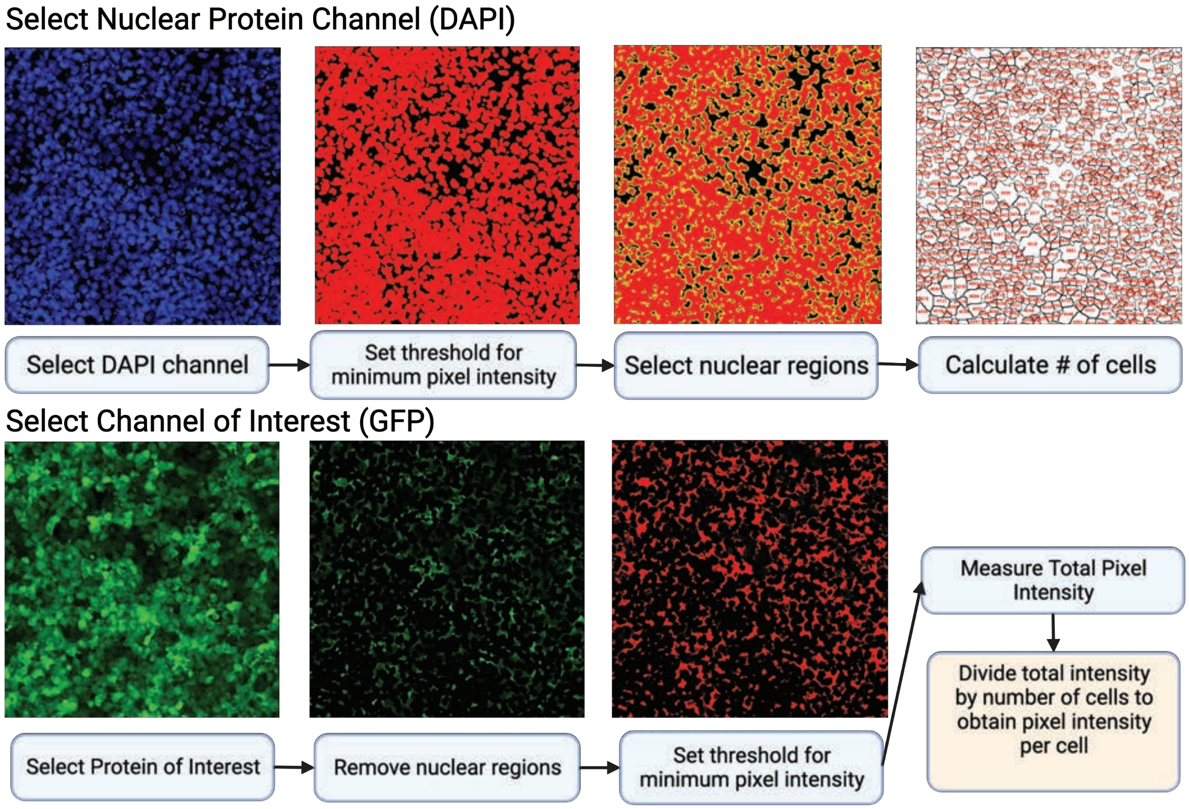


**Supplementary Fig. 3: Graphical schematic of per cell fluorescence intensity quantification using ImageJ FIJI.**

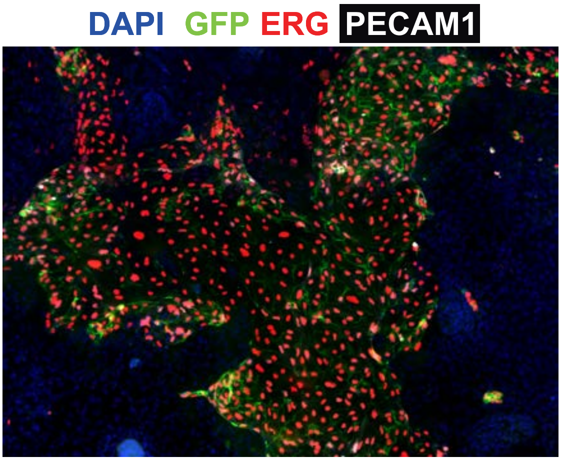


**Supplementary Fig. 4: Immunofluorescence of iETV2-hiPSCs.**

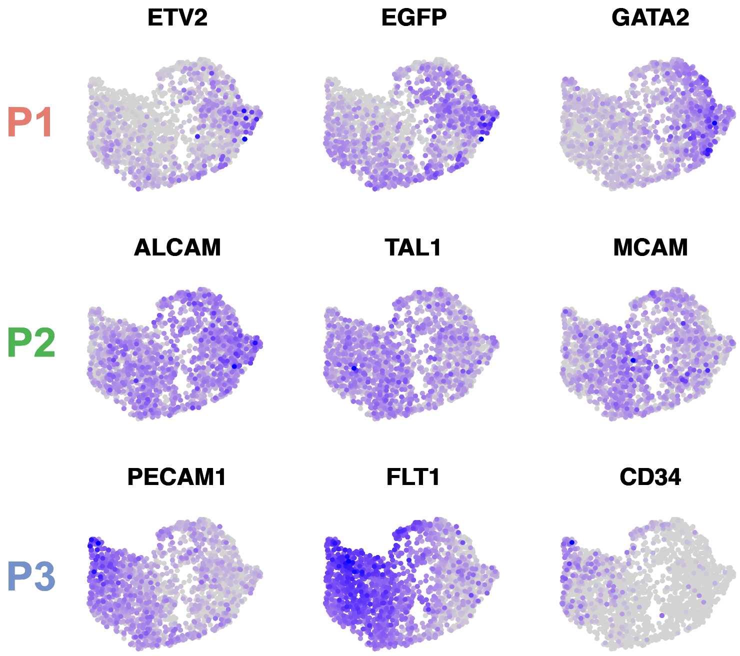


**Supplementary Fig. 5. scRNAseq featureplots of iETV2-hiPSCs.**

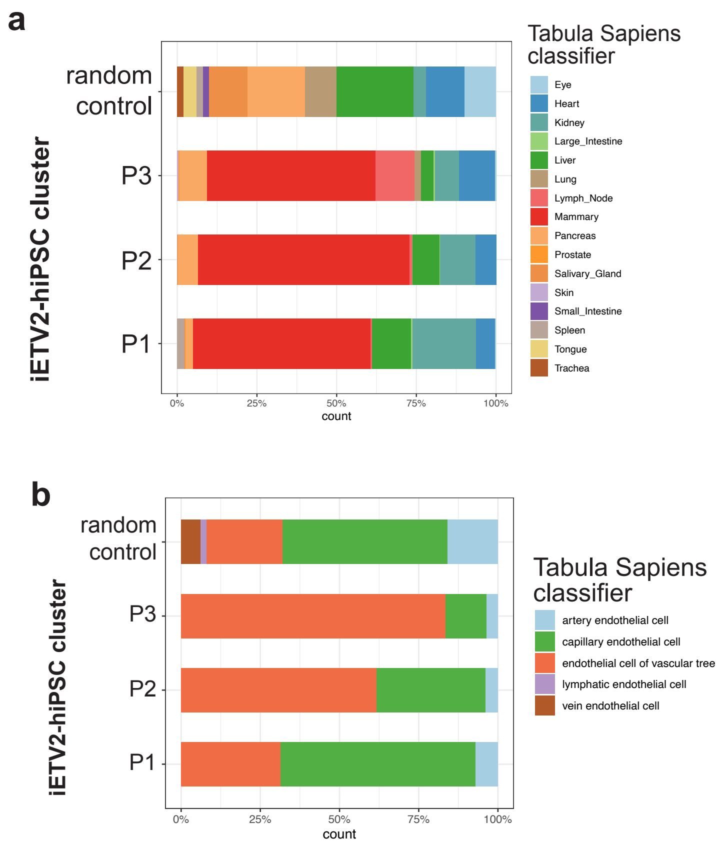


**Supplementary Fig. 6: SingleCellNet Characterization of iETV2-hiPSC scRNAseq.**

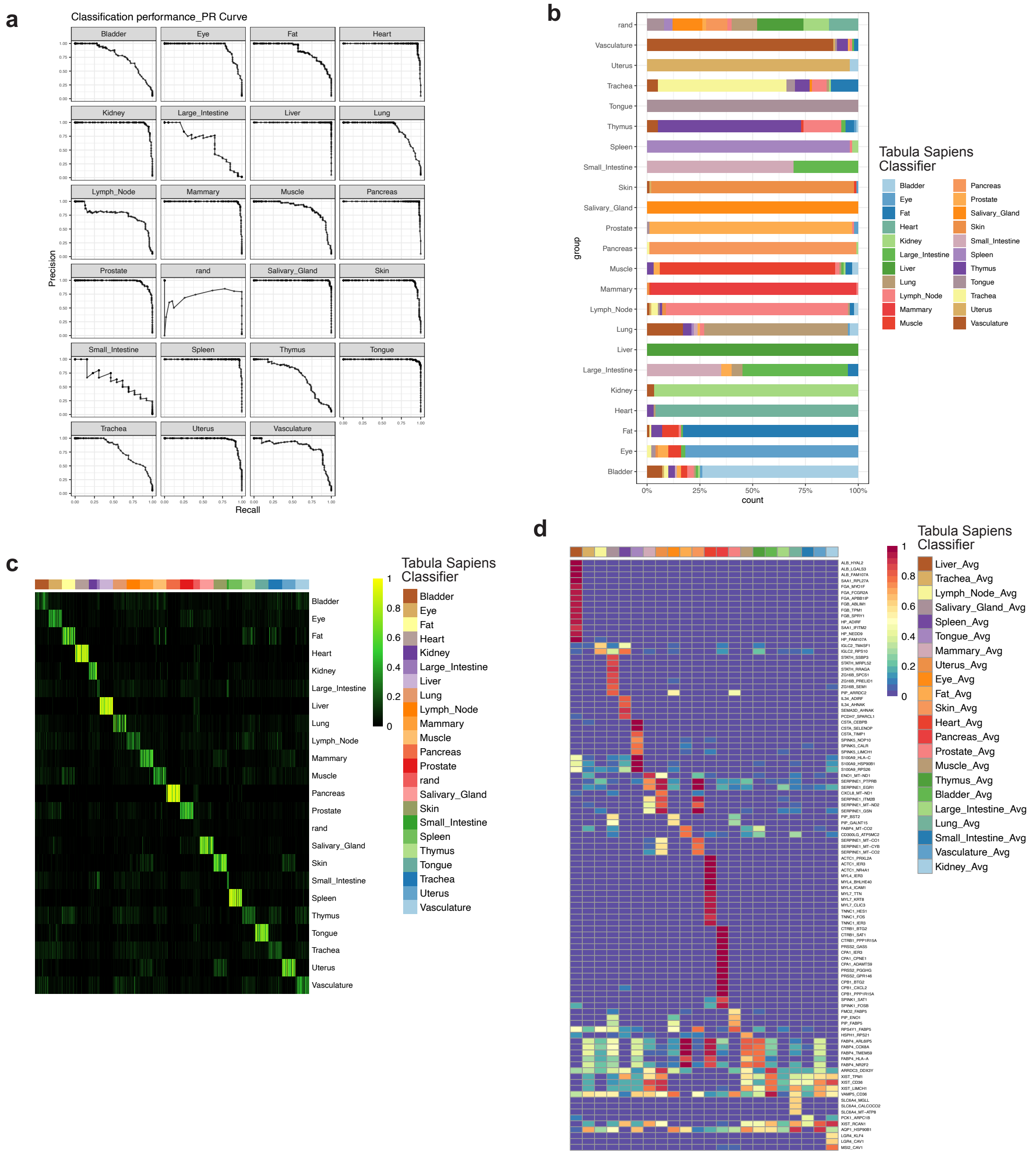


**Supplementary Fig. 7: SingleCellNet using Endothelial Tabula Sapiens by organ system.**

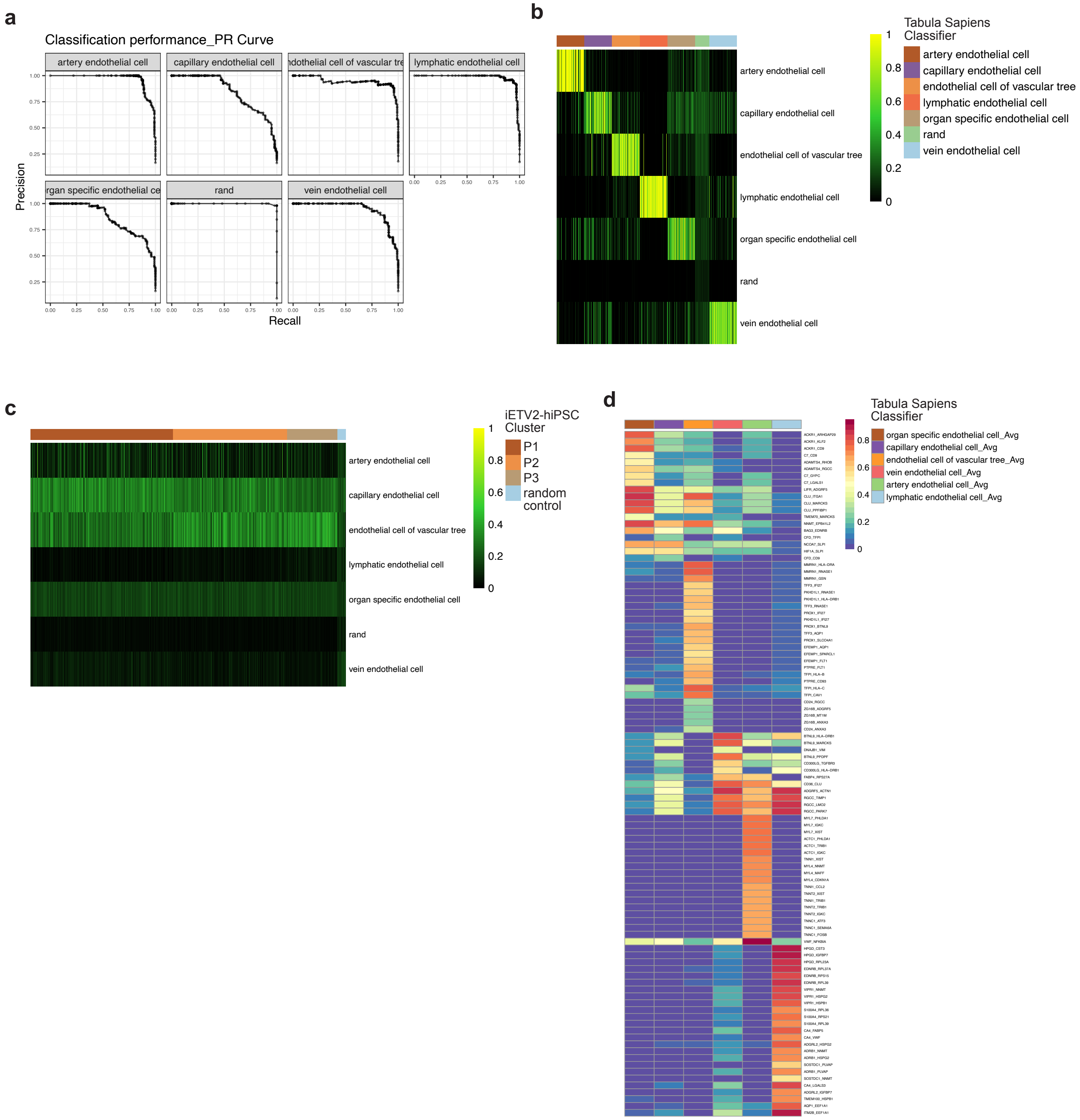


**Supplementary Fig. 8: SingleCellNet using Endothelial Tabula Sapiens Gene Ontology Terms.**

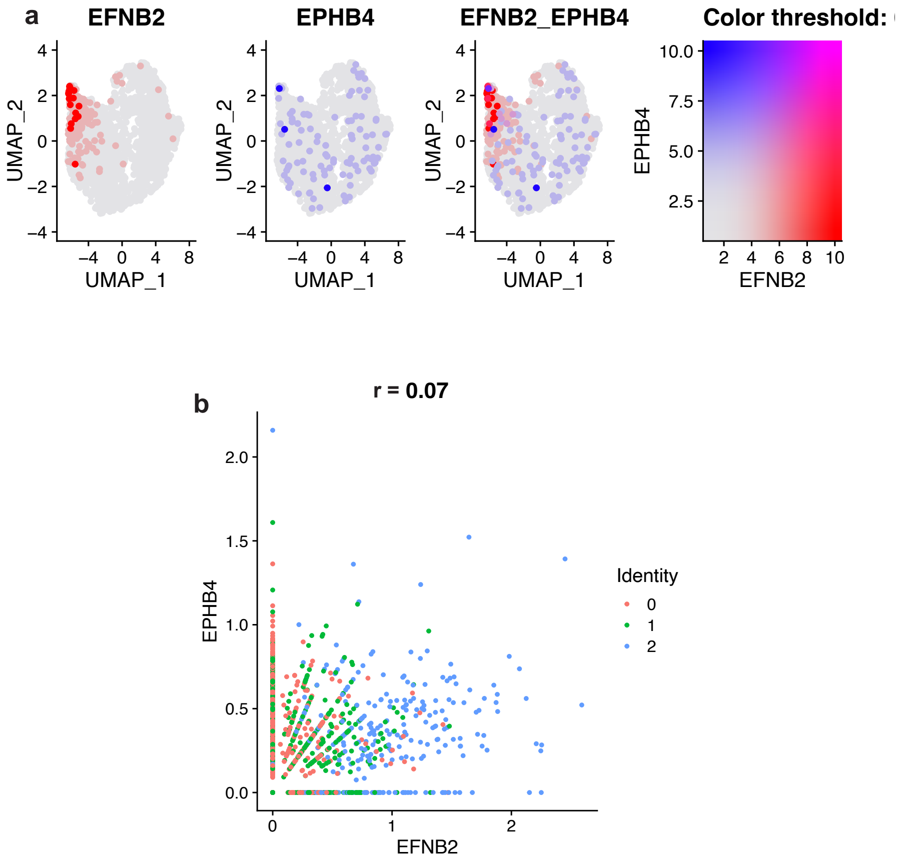


**Supplementary Fig. 9: Arterial and Venous markers of differentiated ETV2 iPSCs.**

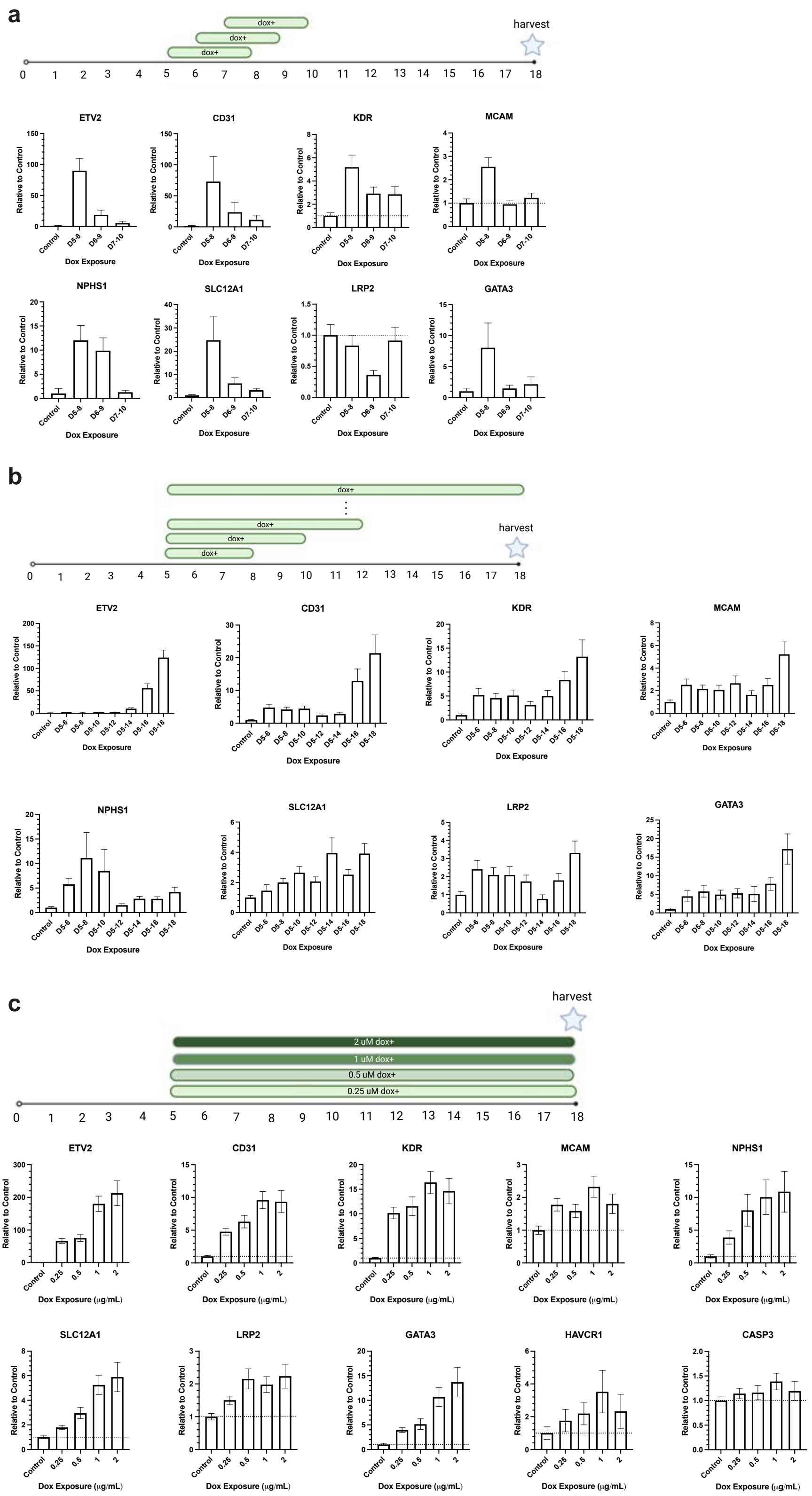

Supplementary Fig. 10: Optimization of doxycycline for vascularization protocol.

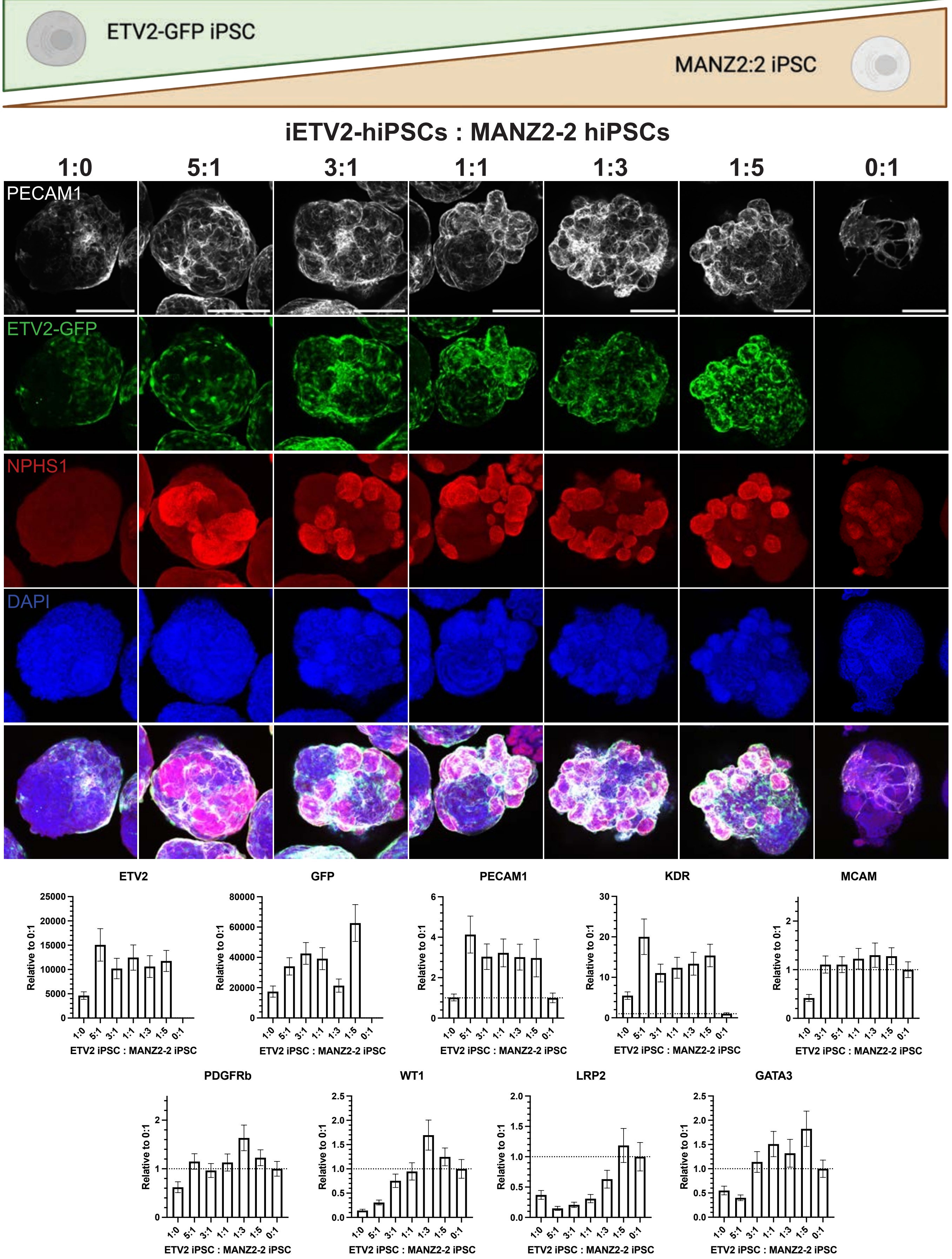


**Supplementary Fig. 11: Optimization of iETV2-hiPSCs to wildtype iPSC ratio for vascularization protocol.**

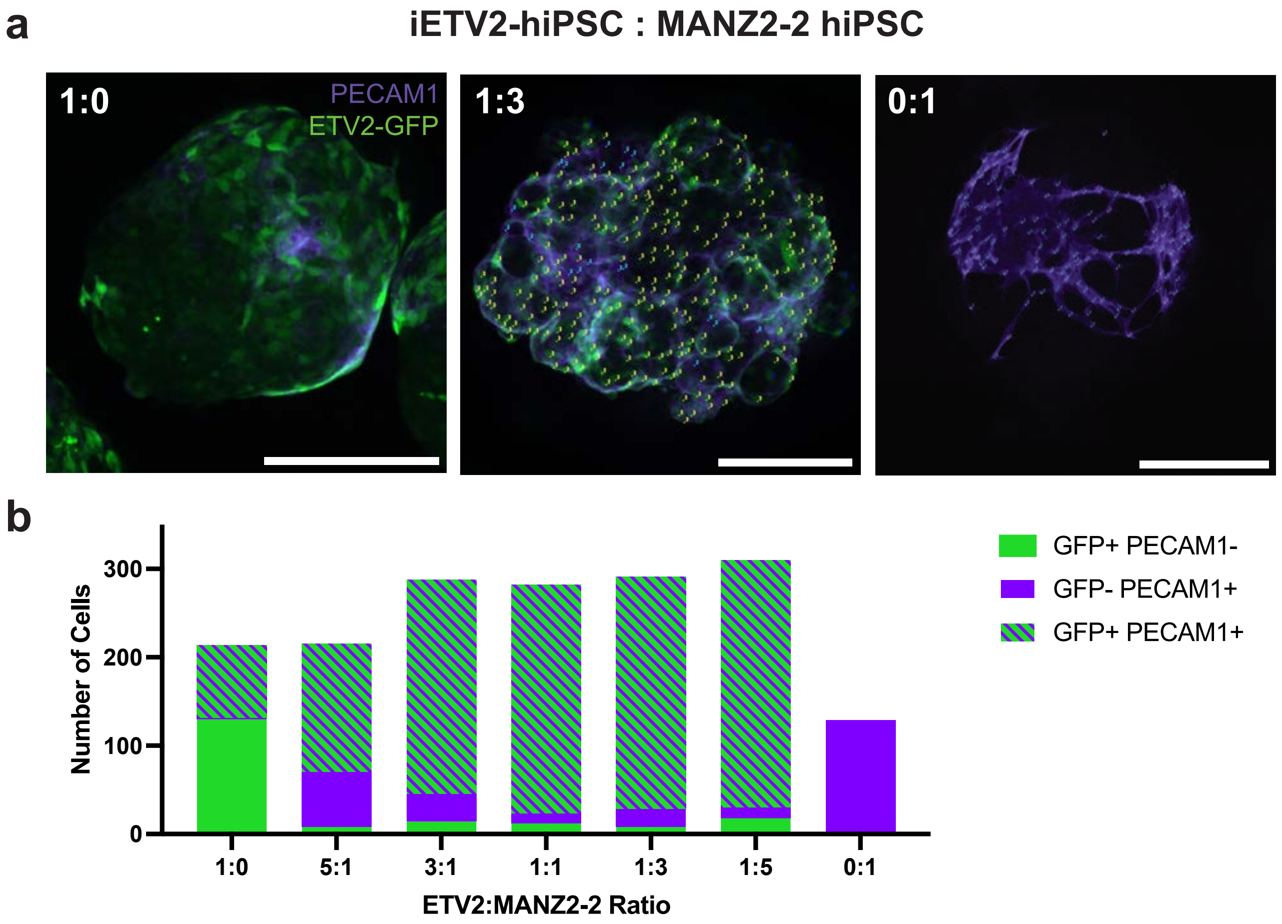


**Supplementary Fig. 12: GFP / PECAM1 colabelling in kidney organoids with iETV2-iPSCs added.**

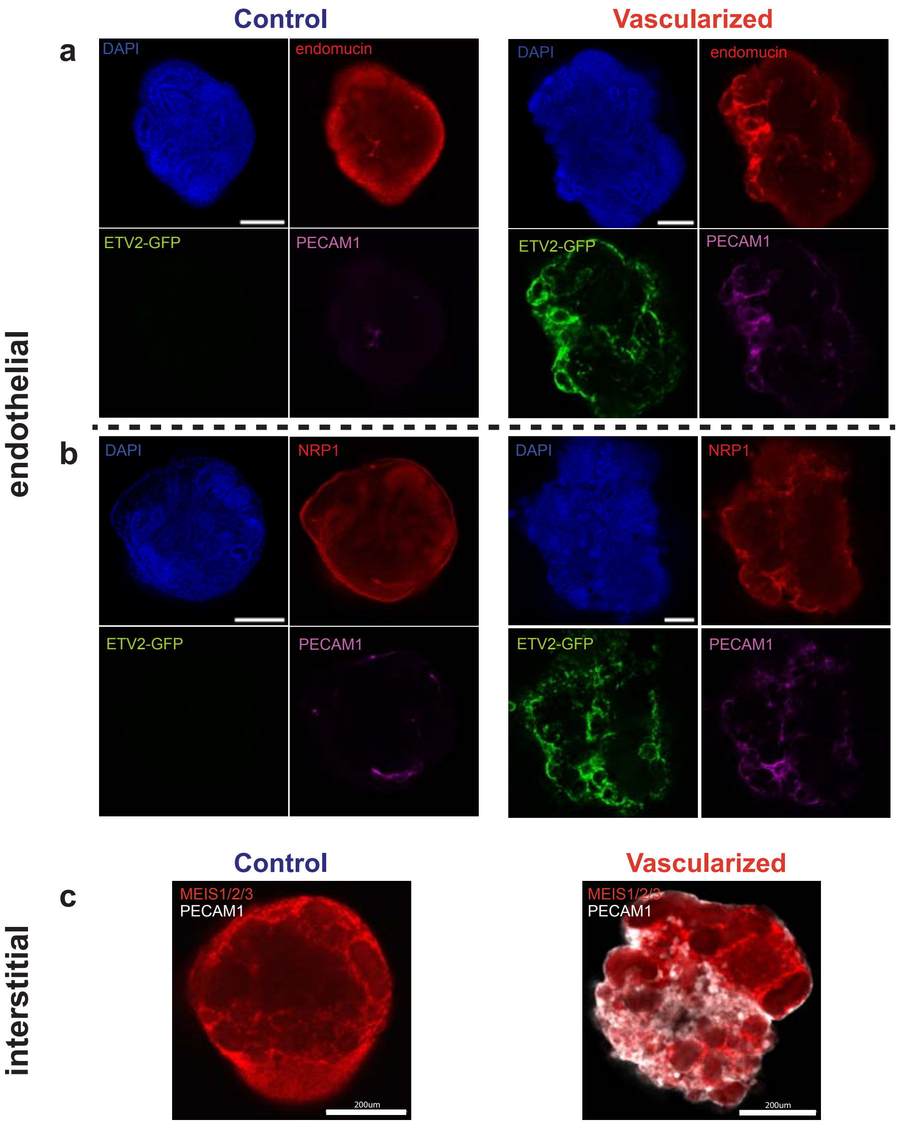


**Supplementary Fig. 13: Control and vascularized kidney organoid immunofluorescent images for endothelia and interstitia.**

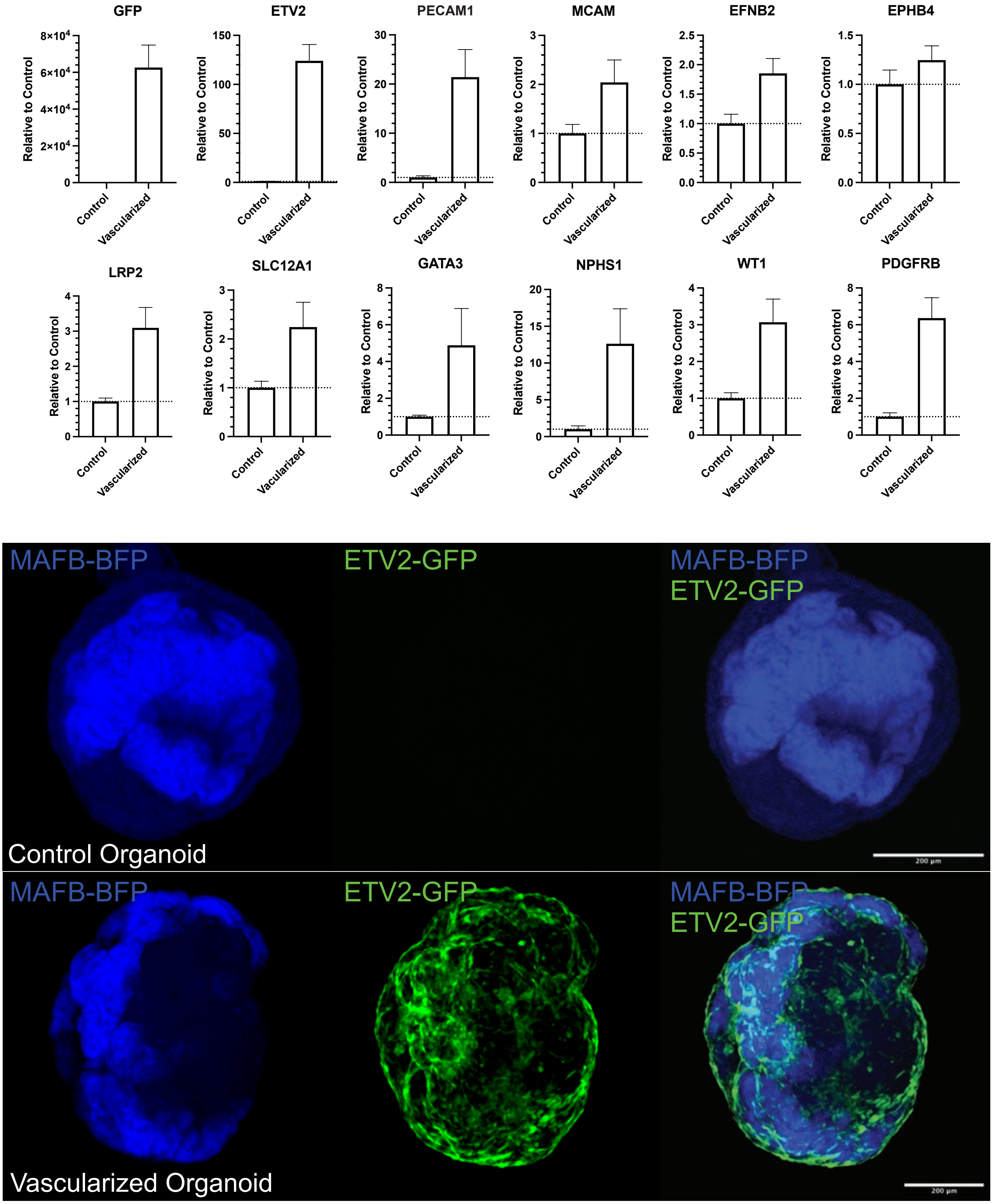


**Supplementary Fig. 14: Triple hiPSC control and vascularized kidney organoids.**

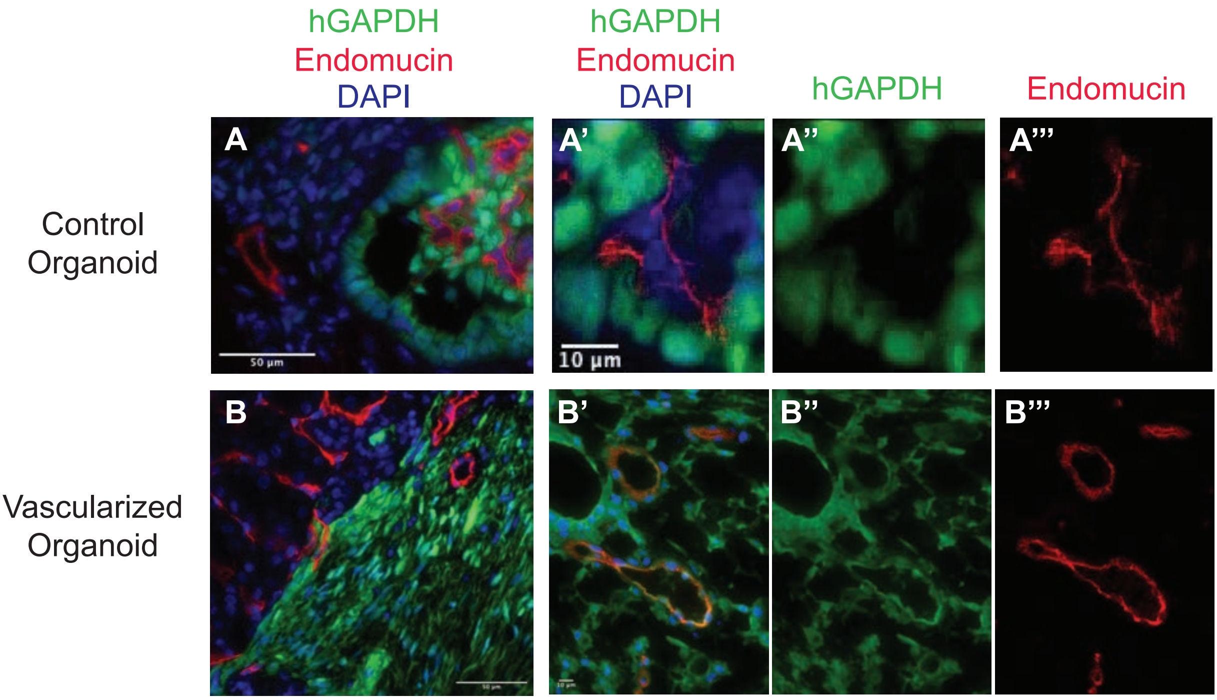


**Supplementary Fig. 15: Control and vascularized kidney organoids implanted under renal capsule of NSG mice.**

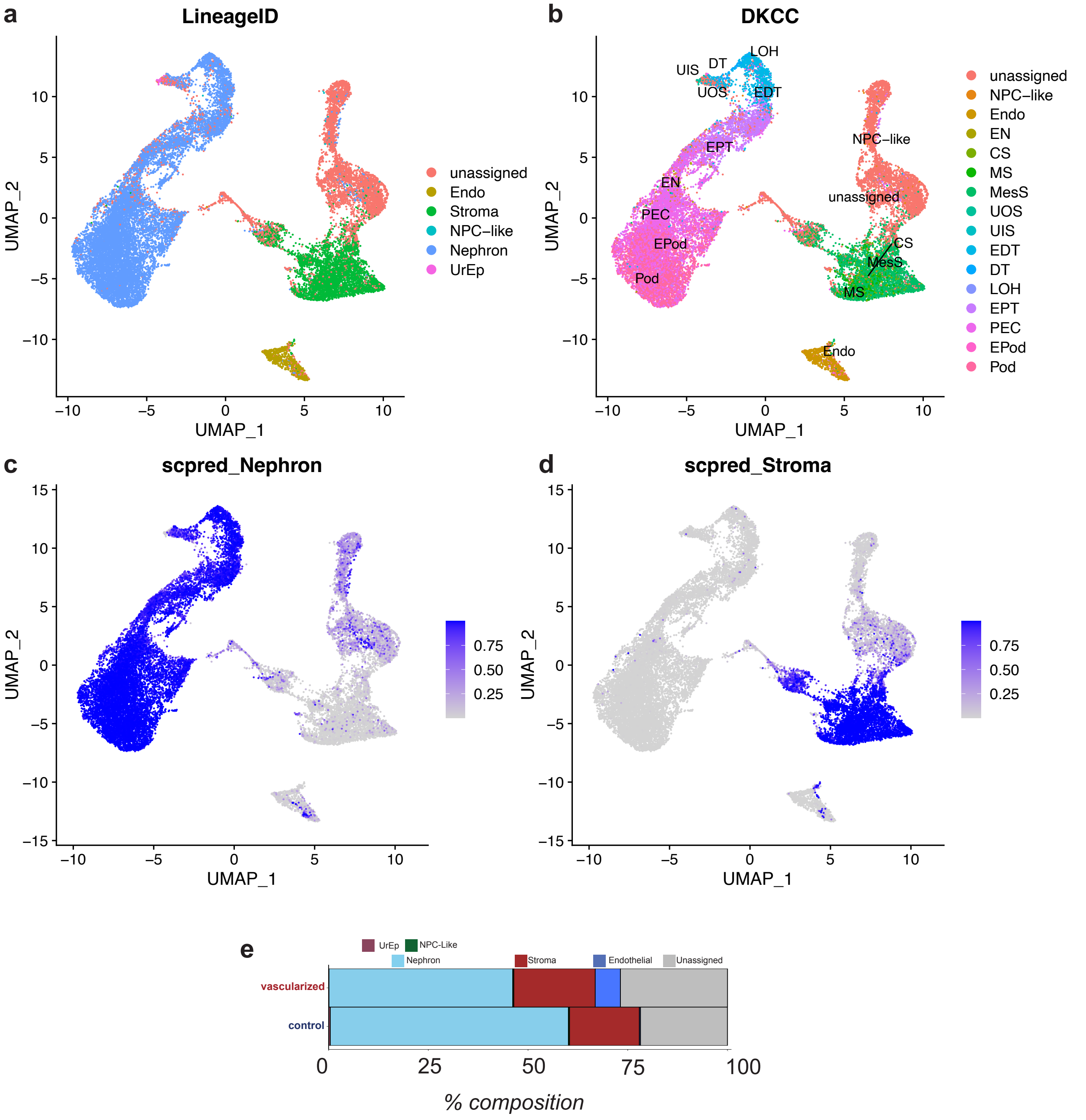


**Supplementary Fig. 16: DevKidCC classification of snRNAseq control and vascularized kidney organoid.**

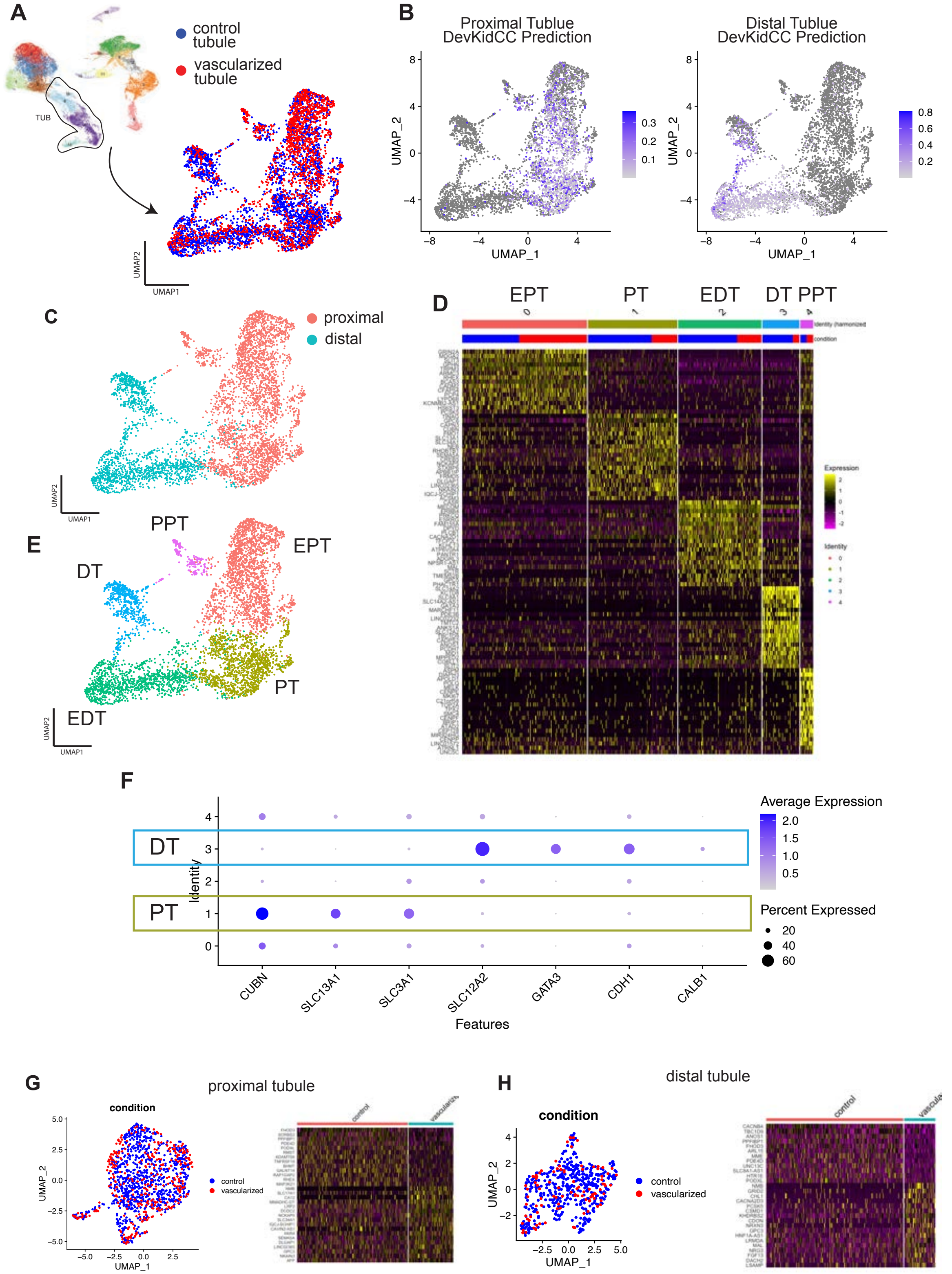


**Supplemental Fig. 17: snRNAseq of control and vascularized tubular cells.**

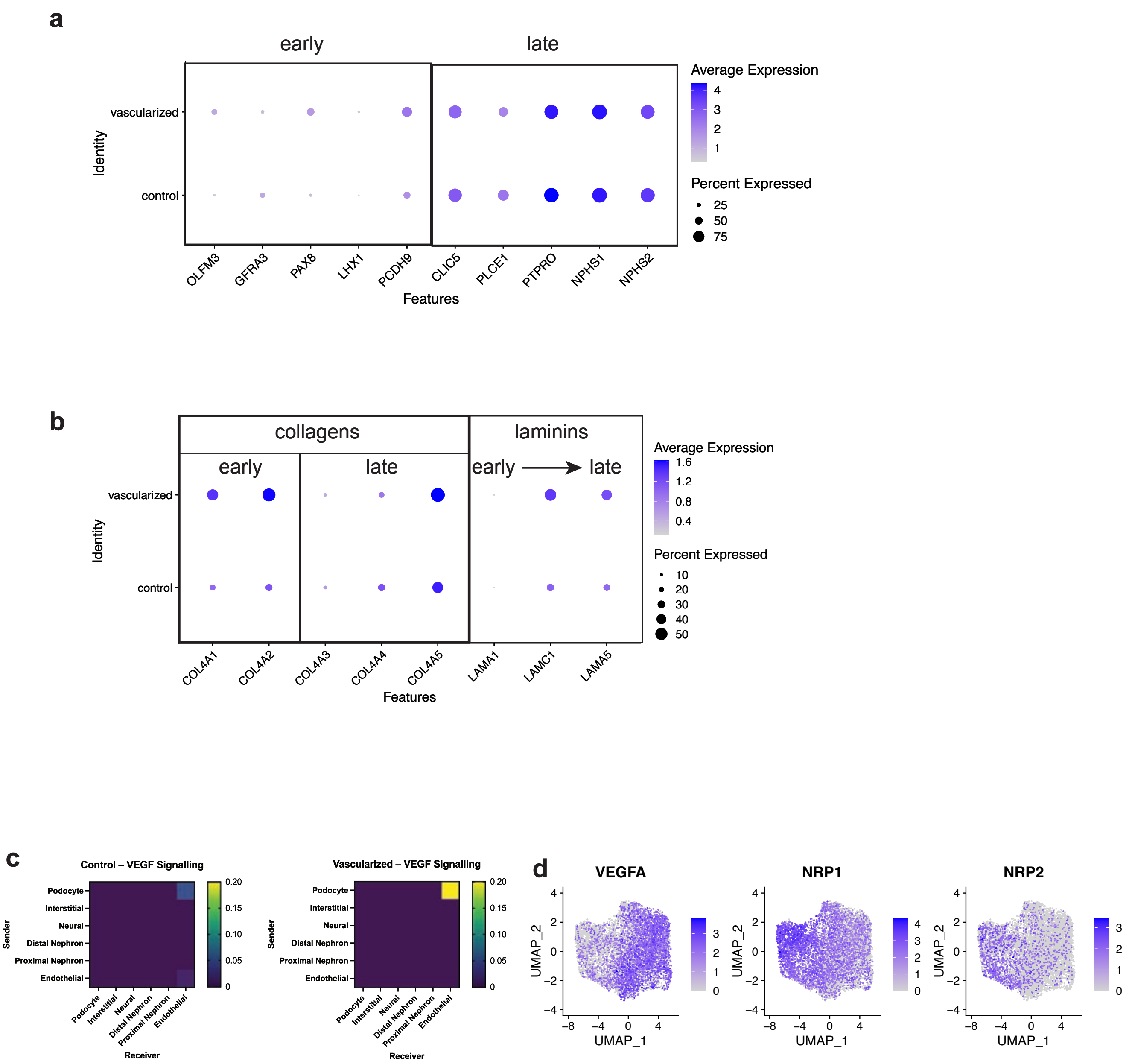


**Supplementary Fig. 18: snRNAseq analysis of podocyte maturation.**

| Category | GO biological process complete | Fold Enrichment | p-value | FDR |
|---|---|---|---|---|
| Basement Membrane | basement membrane organization (GO:0071711) | 8.56 | 5.03E-04 | 2.21E-02 |
| Basement Membrane | regulation of actin filament-based movement (GO:1903115) | 7.9 | 2.03E-04 | 1.06E-02 |
| Basement Membrane | positive regulation of cell-matrix adhesion (GO:0001954) | 7.08 | 3.61E-05 | 2.54E-03 |
| Basement Membrane | regulation of extracellular matrix organization (GO:1903053) | 5.5 | 1.17E-03 | 4.18E-02 |
| Basement Membrane | extracellular matrix organization (GO:0030198) | 3.67 | 1.48E-06 | 1.61E-04 |
| Basement Membrane | extracellular structure organization (GO:0043062) | 3.65 | 1.56E-06 | 1.67E-04 |
| Cell Adhesion | cell adhesion mediated by integrin (GO:0033627) | 10.02 | 3.76E-06 | 3.51E-04 |
| Cell Adhesion | tight junction organization (GO:0120193) | 6.96 | 4.03E-05 | 2.79E-03 |
| Cell Adhesion | regulation of cell-matrix adhesion (GO:0001952) | 6.68 | 1.04E-08 | 1.81E-06 |
| Cell Adhesion | regulation of focal adhesion assembly (GO:0051893) | 6.42 | 6.81E-05 | 4.43E-03 |
| Cell Adhesion | regulation of cell-substrate junction assembly (GO:0090109) | 6.42 | 6.81E-05 | 4.41E-03 |
| Cell Adhesion | tight junction assembly (GO:0120192) | 6.42 | 1.94E-04 | 1.03E-02 |
| Cell Adhesion | bicellular tight junction assembly (GO:0070830) | **6.16** | **6.79E-04** | **2.76E-02** |
| Cell Adhesion | cell-cell junction assembly (GO:0007043) | 6.11 | 8.71E-08 | 1.21E-05 |
| Cell Adhesion | cell-cell junction organization (GO:0045216) | 5.83 | 1.28E-09 | 2.90E-07 |
| Cell Adhesion | cell-matrix adhesion (GO:0007160) | 4.94 | 5.53E-06 | 4.87E-04 |
| Endothelial Maturation | positive regulation of endothelial cell migration (GO:0010595) | 5.76 | 3.02E-06 | 2.98E-04 |
| Endothelial Maturation | endothelial cell differentiation (GO:0045446) | 4.95 | 3.53E-04 | 1.66E-02 |
| Endothelial Maturation | regulation of endothelial cell migration (GO:0010594) | 4.56 | 2.70E-06 | 2.75E-04 |
| Endothelial Maturation | regulation of endothelial cell proliferation (GO:0001936) | 3.69 | 6.00E-04 | 2.55E-02 |
| Endothelial Maturation | positive regulation of angiogenesis (GO:0045766) | 3.38 | 6.51E-04 | 2.69E-02 |
| Endothelial Maturation | positive regulation of vasculature development (GO:1904018) | 3.38 | 6.51E-04 | 2.68E-02 |
| Endothelial Maturation | positive regulation of cell migration (GO:0030335) | 3.38 | 1.24E-09 | 2.87E-07 |
| Endothelial Maturation | regulation of vasculature development (GO:1901342) | 3.17 | 3.21E-05 | 2.31E-03 |
| Endothelial Maturation | angiogenesis (GO:0001525) | 3.14 | 1.31E-05 | 1.07E-03 |
| Endothelial Maturation | blood vessel development (GO:0001568) | 3.12 | 6.25E-08 | 9.24E-06 |
| Glomerular Maturation | positive regulation of glomerular mesangial cell proliferation (GO:0072126) | 22 | 7.52E-04 | 2.95E-02 |
| Glomerular Maturation | regulation of glomerular mesangial cell proliferation (GO:0072124) | 20.54 | 1.13E-04 | 6.68E-03 |
| Glomerular Maturation | regulation of blood volume by renin-angiotensin (GO:0002016) | 17.11 | 1.34E-03 | 4.61E-02 |
| Glomerular Maturation | glomerulus development (GO:0032835) | 5.5 | 1.17E-03 | 4.17E-02 |
| Kidney Differentiation | regulation of metanephric nephron tubule epithelial cell differentiation (GO:0072307) | 22 | 7.52E-04 | 2.96E-02 |
| Kidney Differentiation | positive regulation of epithelial cell differentiation involved in kidney development (GO:2000698) | 22 | 7.52E-04 | 2.95E-02 |
| Kidney Differentiation | pronephros development (GO:0048793) | 19.25 | 1.02E-03 | 3.77E-02 |
| Kidney Differentiation | positive regulation of cell proliferation involved in kidney development (GO:1901724) | 17.11 | 1.34E-03 | 4.60E-02 |
| Kidney Differentiation | regulation of nephron tubule epithelial cell differentiation (GO:0072182) | 15.8 | 2.56E-04 | 1.27E-02 |
| Kidney Differentiation | regulation of cell proliferation involved in kidney development (GO:1901722) | 14.67 | 3.25E-04 | 1.55E-02 |
| Kidney Differentiation | regulation of epithelial cell differentiation involved in kidney development (GO:2000696) | 12.08 | 6.07E-04 | 2.56E-02 |
| Kidney Differentiation | ureteric bud morphogenesis (GO:0060675) | 7.21 | 3.23E-05 | 2.31E-03 |
| Kidney Differentiation | ureteric bud development (GO:0001657) | 6.42 | 3.02E-06 | 2.99E-04 |
| Kidney Differentiation | mesonephric tubule development (GO:0072164) | 6.35 | 3.33E-06 | 3.21E-04 |
| Kidney Differentiation | mesonephric epithelium development (GO:0072163) | 6.35 | 3.33E-06 | 3.19E-04 |
| Kidney Differentiation | renal tubule development (GO:0061326) | 5.9 | 1.66E-05 | 1.30E-03 |
| Kidney Differentiation | nephron tubule development (GO:0072080) | 5.57 | 6.65E-05 | 4.35E-03 |
| Kidney Differentiation | kidney epithelium development (GO:0072073) | 4.87 | 6.41E-06 | 5.52E-04 |
| Kidney Differentiation | renal system development (GO:0072001) | 4.33 | 1.60E-09 | 3.48E-07 |
| Kidney Differentiation | kidney development (GO:0001822) | 4.12 | 1.70E-08 | 2.77E-06 |
| Kidney Differentiation | urogenital system development (GO:0001655) | 3.81 | 1.87E-08 | 3.03E-06 |
| Kidney Differentiation | positive regulation of epithelial cell proliferation (GO:0050679) | 3.68 | 2.88E-05 | 2.11E-03 |
| Kidney Morphogenesis | negative regulation of mesenchymal cell apoptotic process involved in nephron morphogenesis (GO:0072040) | 38.51 | 2.29E-04 | 1.18E-02 |
| Kidney Morphogenesis | regulation of mesenchymal cell apoptotic process involved in nephron morphogenesis (GO:0072039) | 38.51 | 2.29E-04 | 1.18E-02 |
| Kidney Morphogenesis | negative regulation of apoptotic process involved in morphogenesis (GO:1902338) | 25.67 | 5.34E-04 | 2.31E-02 |
| Kidney Morphogenesis | mesonephric tubule morphogenesis (GO:0072171) | 7.08 | 3.61E-05 | 2.53E-03 |
| Kidney Morphogenesis | nephron morphogenesis (GO:0072028) | 7.03 | 4.07E-06 | 3.78E-04 |
| Kidney Morphogenesis | renal tubule morphogenesis (GO:0061333) | 6.42 | 2.40E-05 | 1.81E-03 |
| Kidney Morphogenesis | kidney morphogenesis (GO:0060993) | 6.28 | 3.68E-06 | 3.45E-04 |
| Kidney Morphogenesis | nephron tubule morphogenesis (GO:0072078) | 6.04 | 1.00E-04 | 6.03E-03 |
| Kidney Morphogenesis | morphogenesis of an epithelial sheet (GO:0002011) | 6.04 | 7.47E-04 | 2.96E-02 |
| Kidney Morphogenesis | nephron epithelium morphogenesis (GO:0072088) | 5.87 | 1.21E-04 | 7.07E-03 |
| Kidney Morphogenesis | branching morphogenesis of an epithelial tube (GO:0048754) | 5.25 | 1.27E-06 | 1.41E-04 |
| Kidney Morphogenesis | morphogenesis of a branching epithelium (GO:0061138) | 4.36 | 9.35E-06 | 7.84E-04 |
| Kidney Morphogenesis | morphogenesis of a branching structure (GO:0001763) | 4.11 | 1.73E-05 | 1.35E-03 |

**Supplementary Fig. 19: Podocytes from vascularized kidney organoid predominant cluster gene set enrichment analysis.**

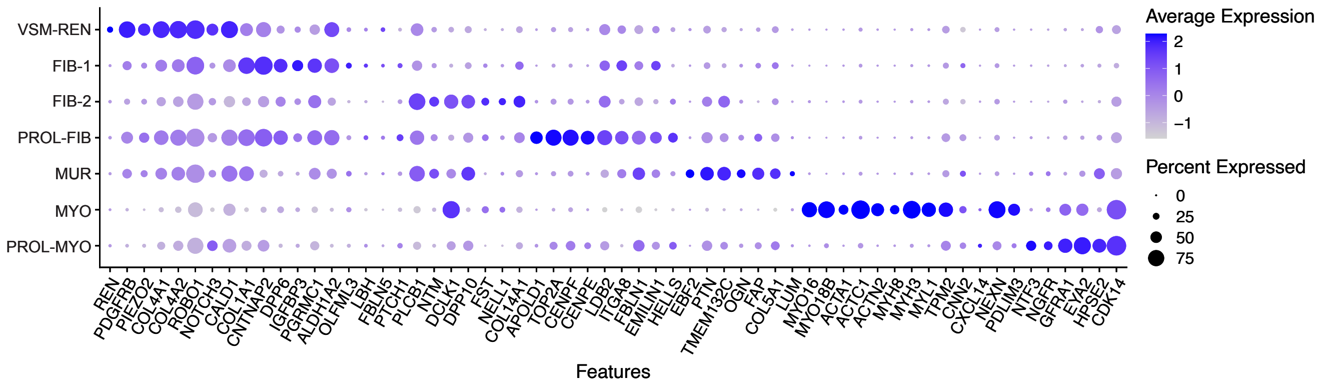


**Supplementary Fig. 20: Canonical genes of interstitial cell populations from interstitial control and vascularized kidney organoid snRNAseq.**

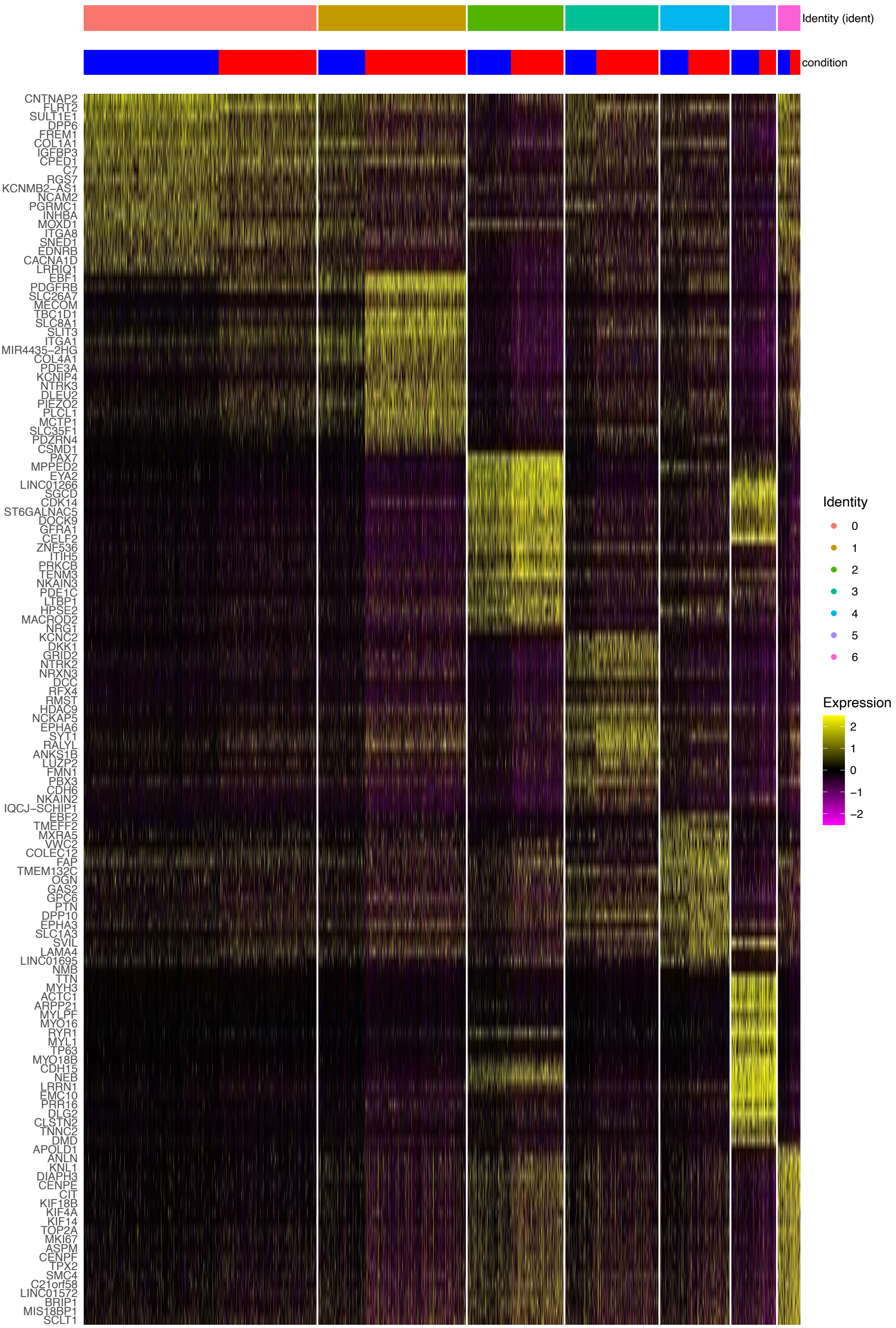

**Supplementary Fig. 21: Differential gene expression in clusters of interstitial cells on snRNAseq.**

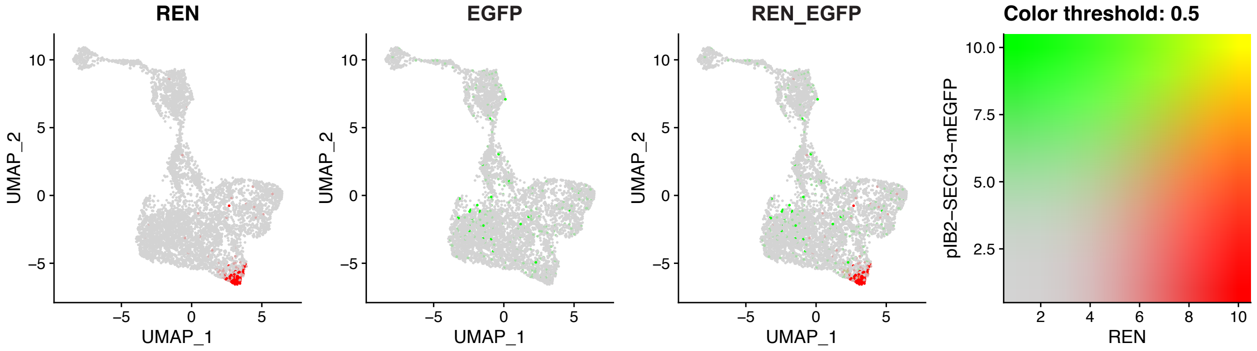


**Supplementary Fig. 22: REN and EGFP expression in control and vascularized snRNAseq interstitia.**

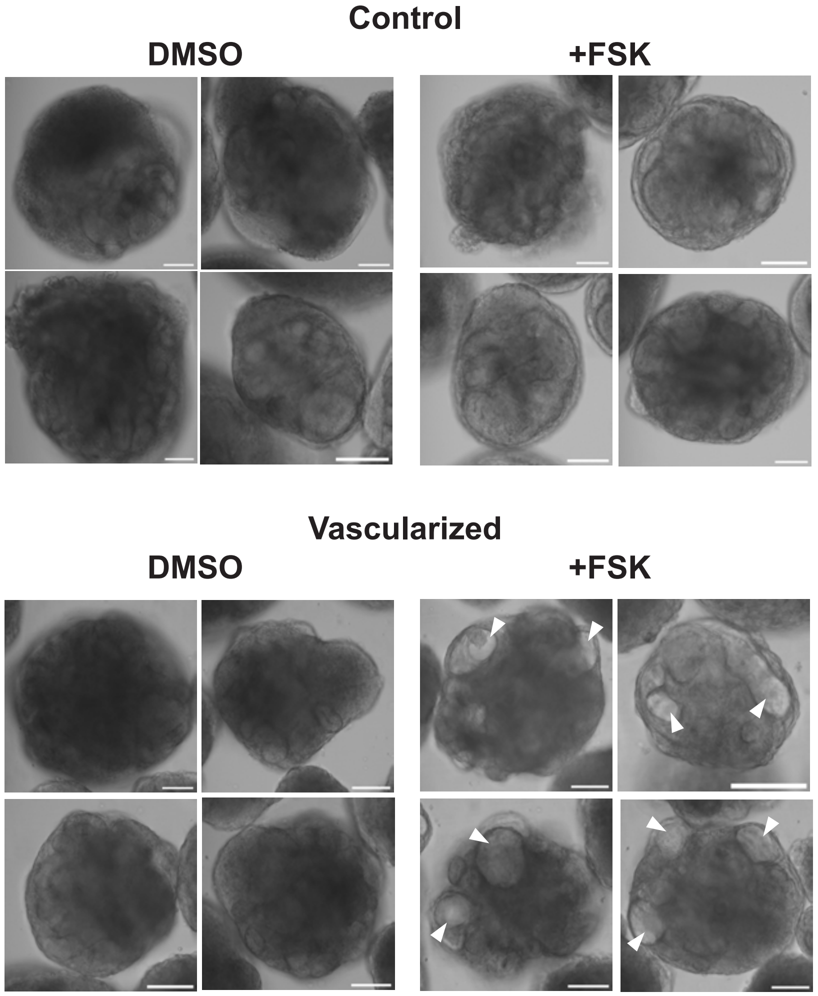


**Supplementary Fig. 23: Light microscope images of forskolin stimulated control and vascularized kidney organoids.**

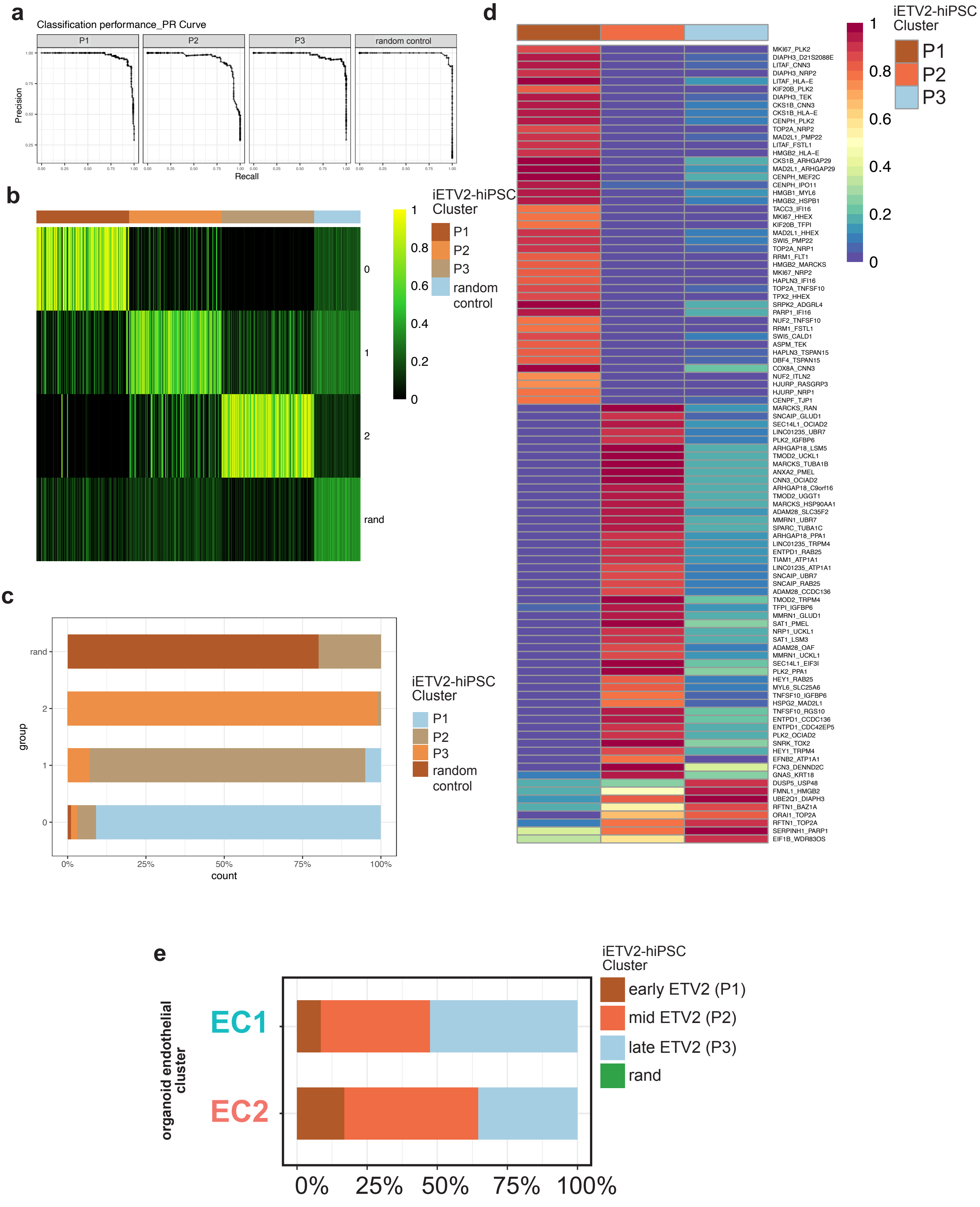


**Supplementary Fig. 24: SingleCellNet training with iETV2-hiPSC scRNAseq dataset.**

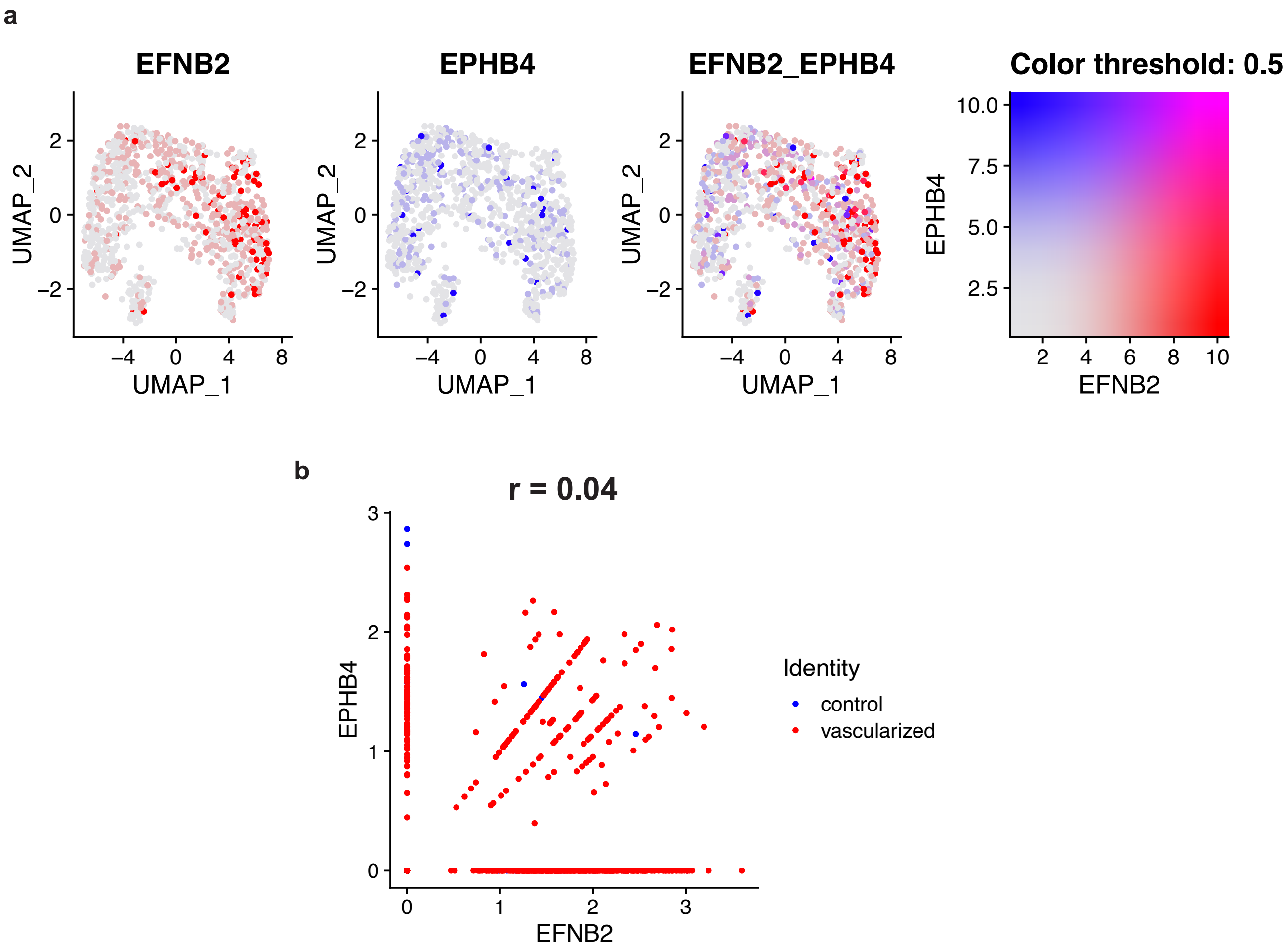


**Supplementary Fig. 25: Ephrin localization in endothelial population of snRNAseq control and vascularized kidney organoid cells.**

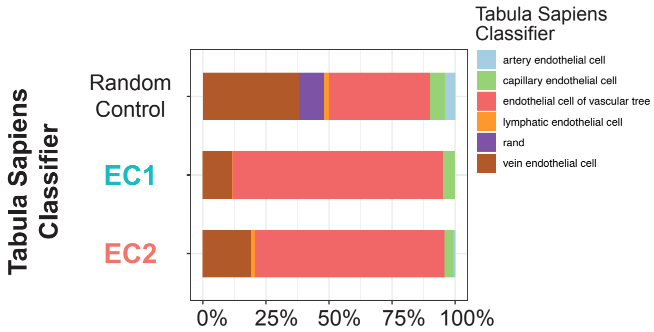


**Supplementary Fig. 26: Morphological classification of endothelial cells in MANZ2-2 control and vascularized kidney organoid using SingleCellNet and Tabula Sapiens endothelial cell gene ontology classifier.**
